# Angiogenic and Immune Predictors of Neoadjuvant Axitinib Response in Renal Cell Carcinoma with Venous Tumour Thrombus

**DOI:** 10.1101/2024.10.22.619358

**Authors:** Rebecca Wray, Hania Paverd, Ines Machado, Johanna Barbieri, Farhana Easita, Abigail R Edwards, Ferdia A Gallagher, Iosif A Mendichovszky, Thomas J Mitchell, Maike de la Roche, Jaqueline D Shields, Stephan Ursprung, Lauren Wallis, Anne Y Warren, Sarah J Welsh, Mireia Crispin-Ortuzar, Grant D Stewart, James O Jones

## Abstract

Venous tumour thrombus (VTT), where the primary tumour invades the renal vein and inferior vena cava, affects 10-15% of renal cell carcinoma (RCC) patients. Curative surgery for VTT is high-risk, but neoadjuvant therapy may improve outcomes. The NAXIVA trial demonstrated a 35% VTT response rate after 8 weeks of neoadjuvant axitinib, a VEGFR-directed therapy. However, understanding non-response is critical for better treatment. We conducted a multiparametric investigation of samples collected during NAXIVA using digital pathology, flow cytometry, plasma cytokine profiling and RNA sequencing. Responders had higher baseline microvessel density and increased induction of VEGF-A and PlGF during treatment. A multi-modal machine learning model integrating features predicted response with an AUC of 0.868, improving to 0.945 when using features from week 3. Key predictive features included plasma CCL17 and IL-12. These findings may guide future treatment strategies for VTT, improving the clinical management of this challenging scenario.

**One Sentence Summary:** A comprehensive multiparametric assessment of the effect of neoadjuvant axitinib in renal cell carcinoma patients with venous tumour thrombus was performed on tissue and peripheral blood, including an integrative machine learning model, which identified both angiogenic and immune determinants of response to therapy.

## INTRODUCTION

Venous tumour thrombus (VTT) occurs in 10-15% of patients with clear cell renal cell carcinoma (ccRCC), where the primary tumour invades the renal vein and inferior vena cava (IVC) and can reach the liver and heart^1,2^. Whilst these patients are technically curable, the surgery required is extensive and complex, requiring multiple teams and the possibility of cardiopulmonary bypass^3,4^. There is considerable morbidity and mortality associated with surgery (5-15%), which increases with the height of the VTT. If left untreated, RCC with VTT has a median survival of 5 months^1,3^.

The NAXIVA trial (Study of Axitinib for Reducing Extent of Venous Tumour Thrombus in Renal Cancer with Venous Invasion, NCT03494816) was a phase II, single-arm, multi-centre study investigating the use of neoadjuvant axitinib, a vascular endothelial growth factor receptor (VEGFR)-directed tyrosine kinase inhibitor (TKI), to reduce the ccRCC VTT. Loss of the tumour suppressor Von Hippel–Lindau (*VHL*) in ccRCC tumours activates the hypoxia response pathway of the cell, leading to induction of VEGF-mediated angiogenesis^5^. VEGFR-TKIs, either alone or in combination with immunotherapy, are used as first line therapy for advanced RCC and have proven efficacy in patients with metastatic disease^6^. In the NAXIVA trial, 35% of patients experienced a reduction in VTT length of >30% after axitinib treatment, leading to less invasive surgery^7,8^. The remaining patients did not benefit from the neoadjuvant treatment. Progress is needed in understanding the reasons for non-response, to improve treatments for these patients.

Little is known about the mechanisms driving treatment response in the VTT. There is evidence that the VTT arises as a rapid outgrowth of the primary tumour and has shared driver events^9^. Studies have shown viable proliferating tumour cells^10^ and immune infiltrate^11,12^ in the VTT. At least one study is currently investigating combination treatment in the specific setting of ccRCC with VTT (NEOPAX, NCT05969496)^13^. However, no prospective study has examined the determinants of VTT response to systemic treatment.

In metastatic ccRCC, RNA-based signatures have been used to group patients according to the therapies most likely to benefit them^14–16^. A prospective study (BIONIKK, NCT02960906) classified patients into four transcriptome-based groups, finding that patients with an immune-low tumour microenvironment (TME) had improved survival during combination of two immunotherapy drugs compared to patients with higher immune-infiltration and inflammatory markers^17^. DNA, protein and clinical markers have also been investigated for patient stratification^18^, but none have been widely adopted for use in metastatic cases, nor in neoadjuvant settings^19–21^. In wider oncology practice, the complexity of the TME and the disparate data generated from multiple sources complicates the development predictive signatures. In other tumour types, the use of machine learning (ML) approaches to integrate data streams has provided valuable insights^22,23^.

To investigate predictive markers of VTT response to axitinib, we performed a comprehensive multiparametric assessment of the TME and peripheral blood for patients in the NAXIVA trial. An ML model was then used to identify biomarkers for VTT response. Identifying reliable predictors of VTT treatment response would allow a personalised approach to treatment selection, which would improve outcomes, avoid overtreatment, and inform the design of future studies for VTT management.

## RESULTS

### Assessment of VTT length response on the NAXIVA Trial

The design and clinical outcomes of the NAXIVA trial have been fully reported elsewhere^7^. Briefly, eligible patients underwent baseline tumour biopsy, then up to 8 weeks of axitinib treatment, followed by surgery to remove the primary tumour and VTT (Fig. 1a). Serial blood samples were collected during the study. Extent of the VTT was assessed by MRI scan at baseline (week 1), week 3 and week 9. For the present study, we included the 20 evaluable patients that were assessed in the trial.

**Figure 1.**
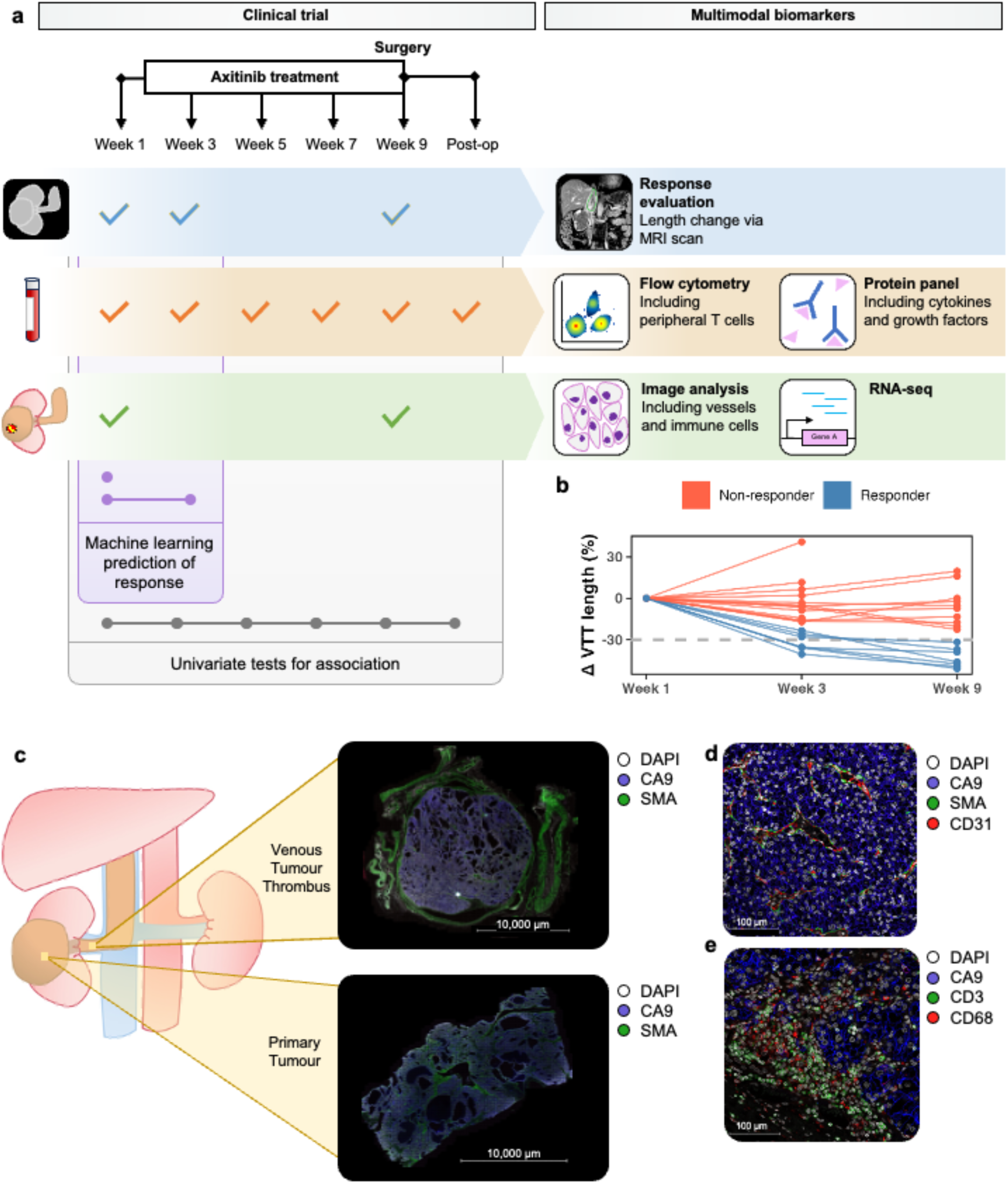
Multiparametric investigation of VTT response in the NAXIVA trial. **a,** Patients received up to 8 weeks of axitinib treatment. VTT response was evaluated by MRI at baseline, week 3 and week 9. Tissue was collected at baseline biopsy, and at surgery from the VTT and primary tumour. Serial blood samples were taken before, during and after treatment. Research samples were assessed by a range of techniques to identify markers of response. Baseline and week 3 parameters were combined in a machine learning model for treatment response. **b,** Patients reaching 30% reduction in VTT length by the end of the treatment course are classed as responders in the NAXIVA trial. 7 of 20 patients were classed as responders. **c,** Whole slide scan of VTT and paired primary tumour, CA9+ viable tumour fills the lumen of the renal vein. **d,** CD31+ microvessels surrounded by SMA+ pericytes are abundant within the VTT TME. **e,** CD3+ T cells and CD68+ macrophages are present within the VTT TME.

In the main trial analysis, the primary endpoint was change in the Mayo level of VTT^7^. However, the Mayo level is a categorical classification based on anatomical landmarks^8^, and relatively small changes in the VTT dimensions may result in a change in Mayo level, or conversely a large change may not cross a Mayo level. To investigate the TME biological response to therapy, we analysed against the continuous percentage change in the VTT length. 7/20 patients achieved a >30% reduction in VTT length by week 9, and we classified this group as responders for our analysis (Fig. 1b, S1a-b). Considering the main clinical parameters, in keeping with the results reported in the clinical study^7^, axitinib dosing, sex and TNM status did not appear to affect the VTT length change after treatment (Fig. S1c-f).

### TME of untreated VTT resembles the primary RCC TME

First, we assessed the microenvironment of untreated resected VTTs, outside of the NAXIVA trial, in comparison to the corresponding primary tumour. In keeping with previous studies, VTT consists mainly of Carbonic Anhydrase 9 (CA9) positive viable tumour cells which fill the vessel lumen (Fig. 1c). In some examples, the morphology of the VTT was very similar to the primary (such as the cystic structures seen in Fig. 1c). There are extensive microvascular structures within VTT, with CD31 positive vessels surrounded by alpha smooth muscle actin (SMA) positive pericytes, between the CA9 positive tumour cells (Fig. 1d). There is immune infiltration of both CD3 positive T-cells and CD68 positive macrophages (Fig. 1e). We assessed the relationship between the VTT TME and corresponding primary tumour by quantitative immunohistochemistry (IHC) for the following key markers in 10 paired cases: Ki67, CD8 and CD31 (Fig. S1g-i). The levels of Ki67, CD8 and CD31 were all significantly correlated between the VTT and primary tumour (CD8 *p* = 0.022, Ki67 *p* = 4 x 10^-4^, CD31 *p* = 0.032, Fig S1g-i). These data demonstrate that the microenvironment of untreated VTT closely resembles that of its parent tumour, and so therapies that are effective against a primary should be effective against VTT.

### Higher microvessel density is associated with VTT response to axitinib

To analyse the effect of axitinib on tumour vasculature, whole slide imaging (WSI) of NAXIVA patients’ baseline biopsy, and post-treatment VTT and primary tumour samples were analysed for microvessel density (MVD) by HALO image analysis (Fig. 2a). The baseline biopsy CD31+/CD34+ MVD in responders was significantly higher than in non-responders (*p* = 7.88 x 10^-4^, Fig. 2b), and was followed by a significant MVD reduction in the VTT after treatment (*p* = 4.06 x 10^-4^). In contrast, the MVD remained at a stable, low level in non-responders (Fig. 2b). This effect was also seen when quantifying CD31+ and CD34+ mono-markers (Fig. S2a-b). Upon assessment of SMA+ cancer associated fibroblast (CAF) area coverage, there was a non-significant trend towards a reduction in the SMA+ CAF in non-responders on treatment (*p* = 0.0867, Fig. S2c).

**Figure 2.**
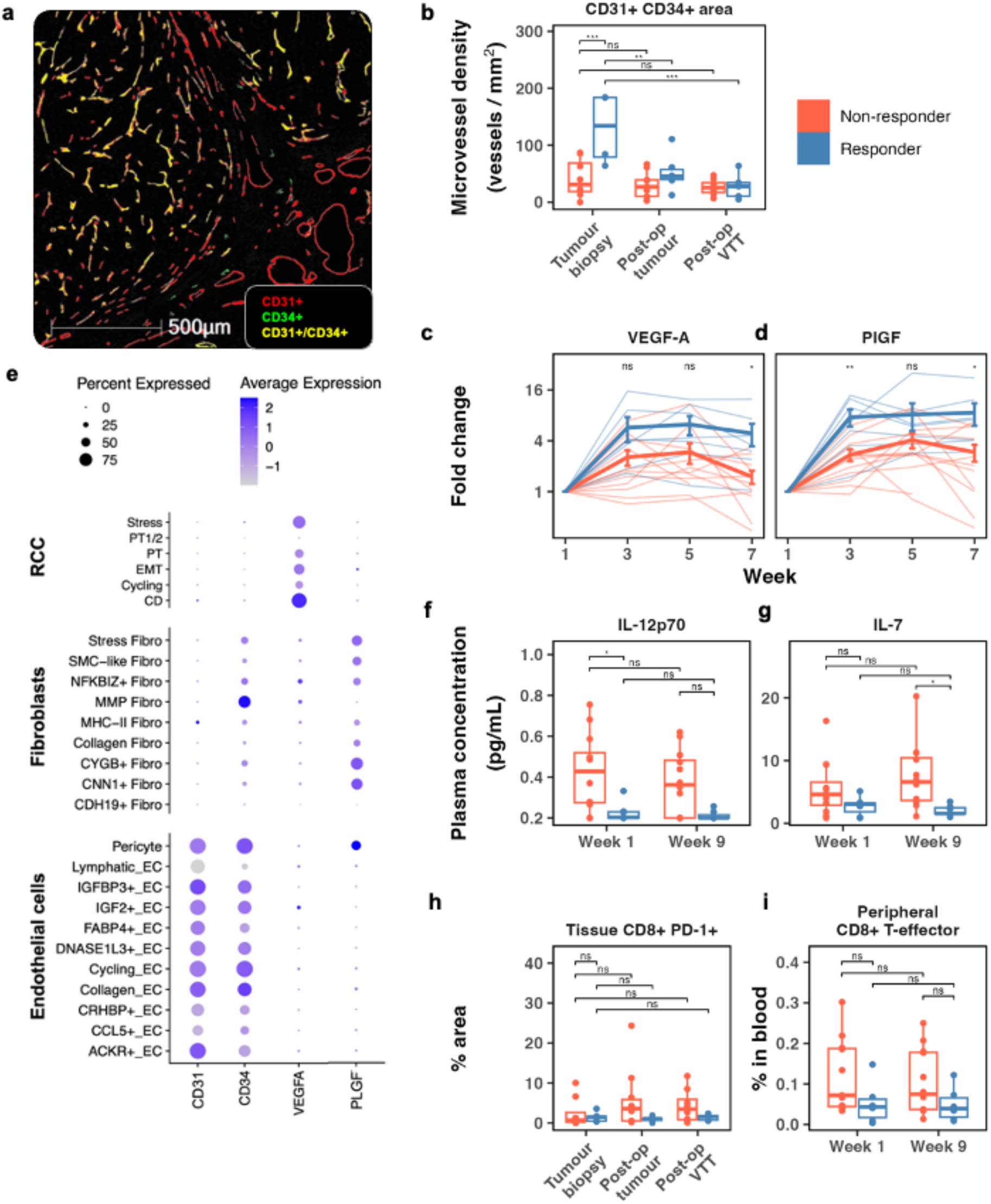
Responder and non-responder phenotypes. **a,** Representative image of HALO analysis markup of microvessels on multiplex immunofluorescence slides. **b,** Responders have higher CD31+/CD34+ microvessel density pre-treatment than non-responders (one-way ANOVA with Tukey’s post-hoc test). **c-d,** Fold change in plasma VEGF-A **(c**) and PlGF **(d**) relative to pre-treatment baseline (thin lines, individuals; bold lines, mean and SEM; unpaired Student’s T-test). **e,** Single cell RNA sequencing analysis of 12 untreated clear cell RCC showing expression of key angiogenesis genes by cell subset. **f-g,** Responders have lower levels of IL-12p70 and IL-7 pre-treatment than non-responders (one-way ANOVA with Tukey’s post-hoc test). **h-i,** Non-responders trend towards higher immune markers in the blood and tissue (one-way ANOVA with Tukey’s post-hoc test). [ns: p>0.05, *: p≤0.05, **: p≤0.01, ***:p ≤0.001].

### Circulating angiogenic factors are differentially induced in responders and non-responders

Axitinib inhibits the signalling response of the VEGF receptors to their soluble ligands; therefore, we assessed the plasma levels of circulating angiogenesis markers during the trial. Absolute levels of VEGF-A were not different either before or during treatment in responders and non-responders (Fig. S3a). However, the fold change relative to each individual patient baseline showed circulating VEGF-A levels increased significantly by the end of treatment in responders compared to non-responders (*p* = 0.0118 at week 7, Fig. 2c). Absolute placental growth factor (PlGF) levels were low at baseline in both groups (Fig. S3b), followed by a strong induction in the responders at week 3 and a return to low levels after treatment ended and the tumour was resected. In fact, there was an approximately 7-fold PlGF increase in responders by week 3 of treatment (*p* = 3.38 x 10^-3^, Fig. 2d). There were some differences in early levels of additional angiogenic markers (Fig. S3), with VEGF-C higher in non-responders at baseline (*p* = 0.0356) and soluble VEGFR1 (sVEGFR1) higher in responders (at week 3, *p* = 0.0445). These markers seemed relatively stable on treatment (Fig. S3).

We then assessed the sources of the key identified angiogenesis markers in a published single cell RNA sequencing dataset of untreated RCC cases^24^, which revealed the primary source of VEGF-A to be the cancer cells; in contrast, PlGF is made by SMA+ myofibroblast subsets and pericytes in the TME (Fig. 2e).

### Non-responders have an immune shift toward CD8+ T cell immunity

Immune features may predict treatment response in advanced RCC^15–17^, we therefore assessed the influence of tissue and blood immune components on VTT response, including the plasma levels of immune cytokines. Circulating IL-12p70 levels were significantly higher in non-responders at baseline (*p* = 0.0282, Fig. 2f). IL-7 levels were significantly higher in non-responders after treatment (*p* = 0.0344, Fig. 2g). There was no difference in interferon gamma, or any other cytokines assessed pre-and post-treatment (Fig. S4).

WSI of biopsy, VTT and primary tumour were analysed by HALO image analysis for T-cell subsets. Comparing responders with non-responders, no significant differences were observed in baseline biopsy CD8+ T-cell levels or CD8+ subsets, including in the CD8+/PD-1+ compartment (Fig. 2h), and this remained stable during treatment in both groups (Fig. S5). No significant differences were seen in overall CD4+ T-cells, or CD4+/Foxp3+ T-regs, or CD68+ macrophages before or after treatment (Fig. S5).

Peripheral blood T-cell subsets were assessed by flow cytometry (Fig. S6a). There was a trend towards increased CD8+ T-cell levels in the peripheral blood of non-responders at baseline (*p* = 0.294, Fig. 2i), and a corresponding shift in the CD4+ to CD8+ T-cell ratio (Fig. S6a). Levels of other CD8+ and CD4+ subsets were similar between groups (Fig. S6a), as were levels of natural killer cells (Fig. S6b) and monocyte subsets (Fig. S6c). There were no differences in B-cell subsets or plasmacytoid dendritic cells (Fig. S7).

### Responders and non-responders have distinct transcriptomic profiles

RNA-seq was performed on pre-treatment biopsies to investigate transcriptomic predictors of response. Principal component analysis of baseline biopsies demonstrated clustering of responders and non-responders (Fig. 3a). Differential gene expression analysis identified some immune-related transcriptomic differences, such as *IL12RB2* (IL-12 receptor beta subunit) and *ARG2* (Arginase type II) (Fig. 3b). However, Gene Ontology (GO) analysis showed that the majority of the most differentially expressed genes are in metabolic pathways, examples including *ALDOB* (Aldolase B) and *ACSBG1* (Acyl-CoA Synthetase, Bubblegum Family, member 1) (Fig. S8). Seven were solute carrier (SLC) family genes, four of which reached high significance (*p* < 0.001). ccRCC survival data from The Cancer Genome Atlas (26) suggests high expression of *SLC6A19* (Sodium-dependent neutral amino acid transporter B(0)AT1)*, SLC22A12* (Solute carrier family 22 [organic anion/cation transporter], member 12) and *SLCO2A1* (Solute carrier organic anion transporter family member 2A1 -a prostaglandin transporter) is associated with improved overall survival, as is *ALDOB* (Fig. S8).

**Figure 3.**
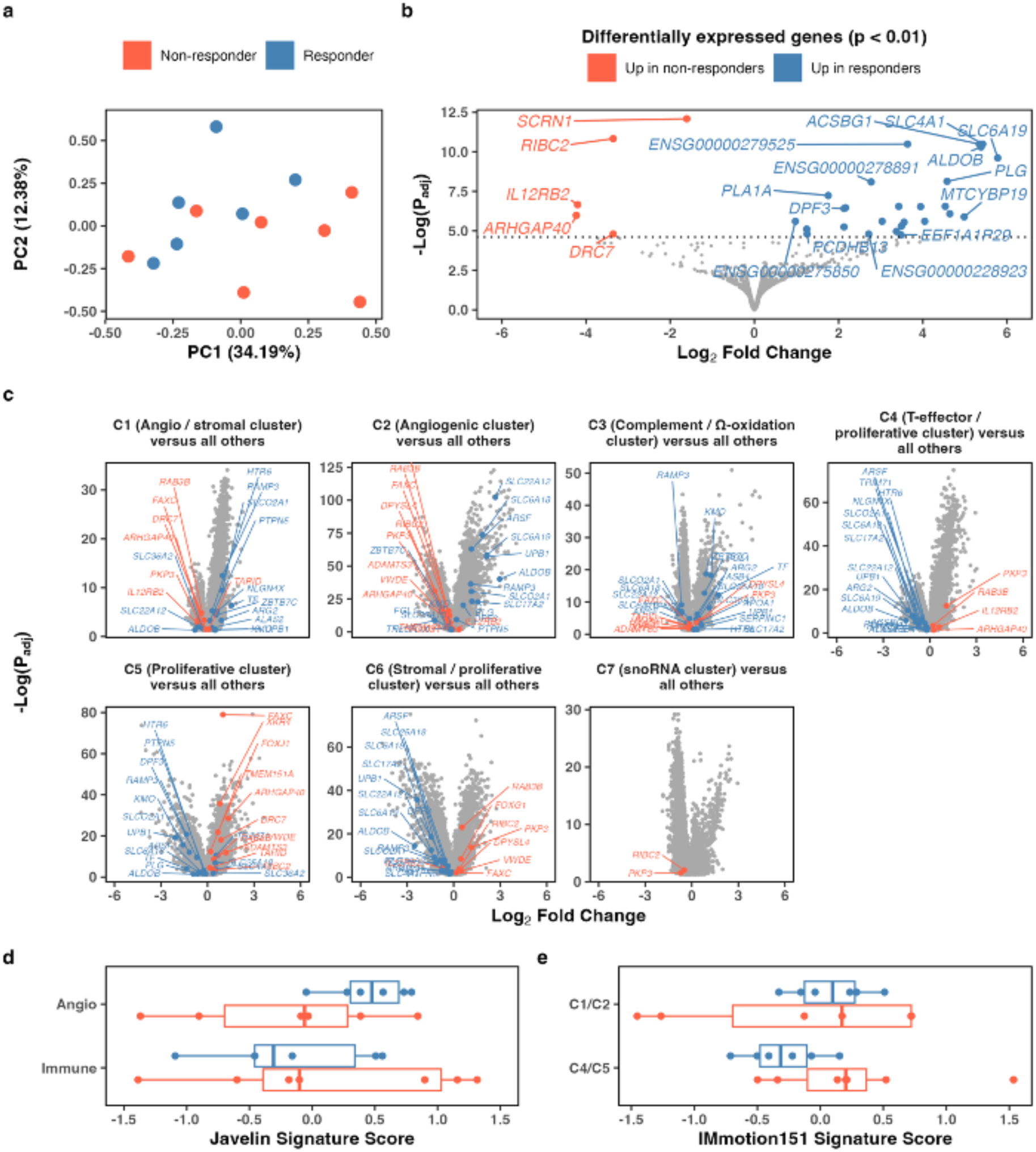
RNA-seq analysis of baseline biopsies. **a,** PCA plot of RNA-seq data for pre-treatment biopsies of responder and non-responder tumours. **b,** RNA-seq results comparing responder to non-responder biopsies via DESeq2. Labelled points are p < 0.01. **c,** Most differentially expressed genes (p < 0.05) plotted on the IMmotion151 RNA-seq clusters [16]. **d,** RNA signature scores for the NAXIVA patients in the transcriptomic signature identified in the Javelin Renal 101 study [15]. **e,** RNA signature scores for the NAXIVA patients in the transcriptomic signature identified in the IMmotion151 study [16].

The most differentially expressed genes from NAXIVA were mapped onto publicly available data from a Phase III study, IMmotion151, which described seven distinct molecular clusters in advanced RCC^16^. Genes highly expressed in NAXIVA responders were also highly expressed in IMmotion151 clusters C1 (angiogenesis/stromal) and C2 (angiogenesis), including the *SLC* family members. In contrast, patients in clusters C4 (T-effector/proliferative), C5 (proliferative) and C6 (stromal/proliferative) have lower expression of the genes highly expressed in NAXIVA responders, and higher expression of the genes highly expressed in NAXIVA non-responders (Fig. 3c).

Published RNA-based predictive signatures of RCC treatment response to anti-angiogenic or immunotherapy from IMmotion151 and Javelin Renal 101, another large phase III clinical trial, were used to calculate scores for each patient in NAXIVA (Fig. 3d-e). NAXIVA responders score higher in the Javelin Renal 101 “Angio” RNA signature^15^, on average. Three of the non-responders score highly in the Javelin “Immuno” score, but the spread of the scores is broad (Fig. 3d). The IMmotion151 molecular subset clusters used to differentiate patients in the ongoing Phase II OPTIC-RCC study^16,25^ – C1/2 angio/stromal, and C4/5 T-effector/proliferative – again showed correlation with the NAXIVA responder and non-responder groups, with non-responders achieving a higher C4/5 score on average than responders (Fig. 3e). However, the C1/2 score is less able to differentiate between these patients.

### A machine learning model integrating multiple baseline features predicts treatment response

Integrating multiple data strands into a predictive model may provide better insights into the drivers of response in oncology trials^22,23^. We developed an ML approach using baseline (week 1) features to predict response outcome, considering a binary classification of response as above, where response is defined as a >30% reduction in VTT length compared to baseline. The input data consisted of 62 features measured for each of the 20 patients. To reduce overfitting of the model to the small dataset, highly correlated features were reduced as part of data pre-processing, and the first part of the model involves a dimensionality reduction step, which selects the features contributing most to response (Fig. 4a). In this model, three features of the 62 were selected. A logistic regression was then fitted to the reduced dataset. We used a leave-one-out cross-validation approach, whereby the feature selection and model training was repeated for each group of 19 patients, generating 20 models which each predicted the response of the remaining one patient. While the number of patients in the study is relatively low, this approach prioritises feature identification for further investigation and follow up in future studies.

**Figure 4.**
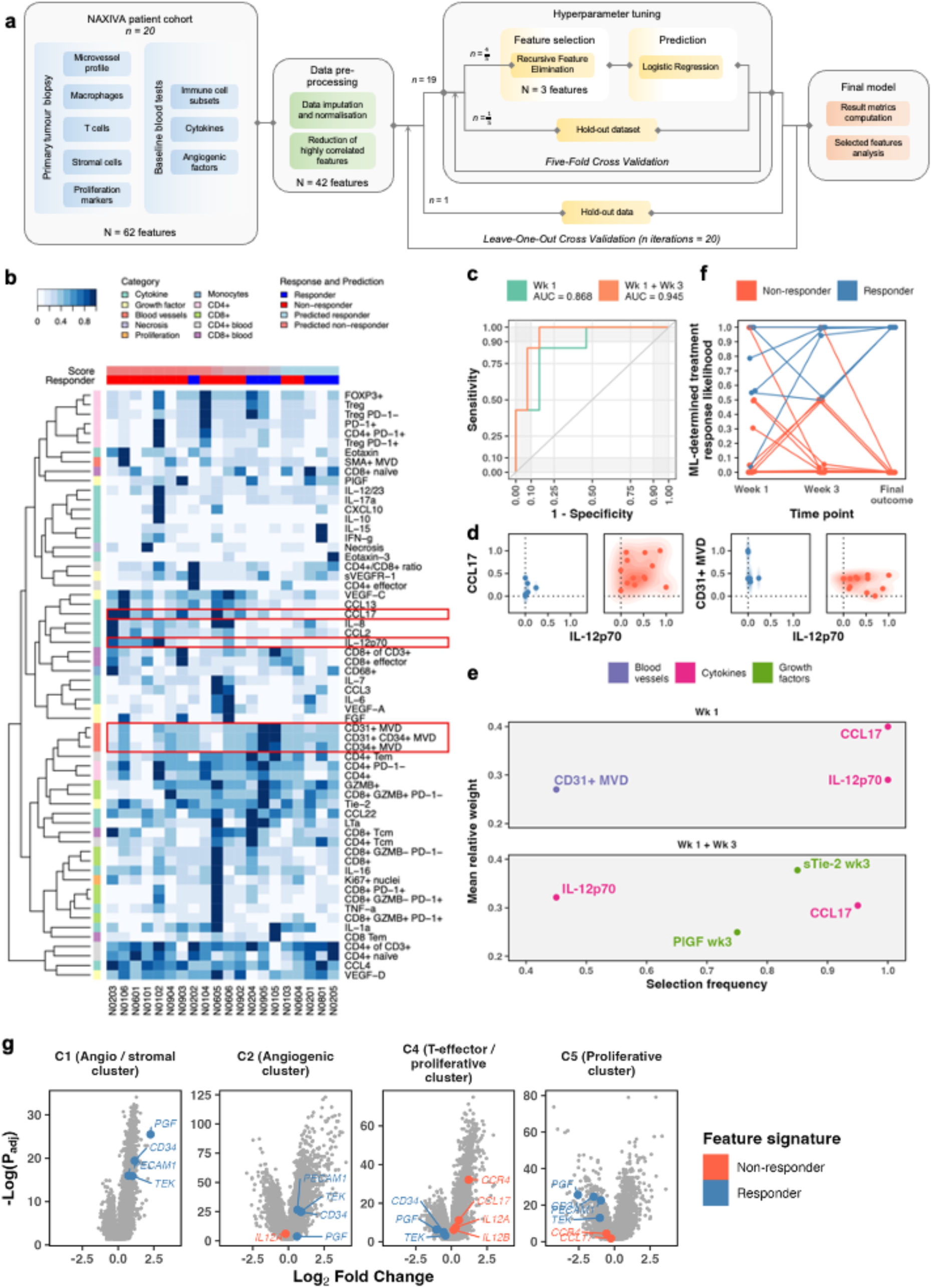
Machine learning model predicts response to axitinib. **a,** Machine learning model workflow. **b,** Pre-processed data description and model-predicted scores for each patient. **c,** Receiver operating characteristic curve. **d,** Selection frequency for selected features (the number of times the feature was selected across the leave-one-out cross-validation iterations divided by the total number of iterations) and mean relative weight of features selected in more than 40% of the iterations. **e,** Density plots of scaled values of two features with highest selection frequency for responders and non-responders. **f,** Prediction of response increases in accuracy and confidence when Week 3 measurements are included in the analysis. **g,** Signature from NAXIVA blood data displayed on IMmotion151 RNA-seq data [16].

The scaled data input to the model are shown in Fig. 4b. With these baseline features, the model achieved an area under the receiver operating characteristic curve (AUC) of 0.868 (Fig. 4c, Table 1). Three specific features were selected repeatedly in at least 8 of the 20 iterations of the model (Table 2). These were found to be plasma IL-12p70, CCL17, and microvessel density. CCL17 was not identified in the univariate data analysis (Fig. S4) but had the highest selection frequency and relative weight assigned by the logistic regression. Responders were always low in circulating CCL17 and IL-12p70, and generally, but not exclusively, higher in tumour MVD (Fig. 4d). Published scRNA-seq data shows that CCL17 is not expressed by RCC cells but may be expressed by conventional dendritic cells in the TME (Fig. S9). The CCL17 receptor, CCR4, is expressed on CD4+ T-cells and highest on CD4+ T-reg cells. IL-12 receptors are widely expressed on T-cell and NK populations (Fig S9).

### Adding early dynamic measure improves the performance of the machine learning model

A challenge in biomarker development for all cancers is the inherent variability between patients, either due to tumour differences or their underlying physiology. Early markers of response after treatment has begun may be informative for clinical decision making. Therefore, we updated the model to include measurements of fold-changes in the plasma angiogenic factors after three weeks of axitinib treatment (Fig. S10). The model achieved a higher AUC of 0.945 (Fig. 4c), with high selection of CCL17 and IL-12p70, as before (Table 2, Fig. 4e). Interestingly, the week 3 fold-change in plasma sTie-2 and PlGF also showed potential for stratification.

Comparing the longitudinal performance of the models, the dynamic model returns higher confidence in the responder classification, which we interpret as a higher probability of response, giving a score of >0.5 for all seven responders (and five of them >0.9). The baseline model gives six responders >0.5 (and three of them >0.9), with one misclassified as <0.1 (Fig. 4f). The selected features correspond as expected to the molecular clusters in the IMmotion151 RNA-seq data (Fig. 4g). PlGF is much higher in the angio-stromal cluster, C1, in keeping with its stromal origin in the single cell analysis.

## DISCUSSION

Our study draws on a unique sample set from a phase II clinical trial, with tissue and serial blood samples taken before, during and after treatment. We conduct a multiparametric analysis including tissue factors by digital pathology and RNA-seq, and both cell-based and soluble-factor analysis of peripheral blood. Furthermore, we deploy ML approaches to prioritise and integrate the parameters. This allows us to gain new insights into determinants of response in ccRCC with VTT, a challenging clinical scenario.

The highly organised nature of the VTT TME is striking, with an established microvessel network, stroma, and immune infiltrate as seen in primary ccRCC. High baseline MVD was predictive of response to neoadjuvant axitinib, particularly in the CD31/CD34+ subset of vessels. Good response was also associated with greater induction of angiogenic growth factors, particularly stroma-derived PlGF. Non-responders had an immune-high phenotype, with higher levels of IL-7 and IL-12, and trends to increased circulating CD8+ T-effectors in blood and CD8+/PD-1+ T-cells in the TME. Assessing transcriptomic data from baseline biopsies, several genes were associated with good response, notably in the solute carrier gene family. The ML model selected IL-12p70, CCL17 and biopsy MVD for response prediction, with model performance improvements seen after the inclusion of early response data, selecting for sTie-2 and PlGF induction. The identified features could be readily assayed in clinical practice.

Our data supports previous reports that the VTT is closely related to the parent tumour and is essentially primary tumour existing within the lumen of the vessel^10–12^. Our finding that VTT axitinib responders have a pro-angiogenic, immune-low phenotype is in keeping with observations in the metastatic setting, where an angiogenesis-rich subgroup is proposed to benefit from VEGFR-TKI therapy^14–17^. Amongst circulating factors, PlGF has previously been described as a pharmacodynamic marker for TKI treatment^26,27^; however, it has not previously been found to be a predictive marker for therapy outcome as demonstrated here. IL-7 supports lymphocyte proliferation^28^, IL-12 is critical for cytotoxic T-cell differentiation^29^, and there is evidence these cytokines may cooperate to enhance antitumour immunity^30^.

Our study finds that highly upregulated genes in NAXIVA responders were also upregulated in C1/2 patients (angiogenic/angio-stromal) in publicly available IMmotion151 transcriptomic data, whereas the opposite was observed for C4/5/6 (T-effector/proliferative, proliferative and stromal/proliferative), which fitted better with the non-responding patients. When published RNA signatures were applied to the NAXIVA transcriptomic data, we found some correlation with outcome, particularly for the Javelin Renal 101 ‘Angio’ score.

The RNA-seq data from the patient biopsies revealed an association between response to axitinib and a number of genes involved in several metabolic pathways, including genes in the solute carrier family, namely *SLC6A19, SLC22A12, SLCO2A1* and *SLC4A1*. The concomitant increase in MVD and expression of genes related to solute metabolism and transport point towards a relationship between the metabolic pathways in the tumour and the induction of angiogenesis. For example, *SLCO2A1* (a prostaglandin transporter) may regulate the endothelial response to prostaglandins^31^, influencing angiogenesis and potentially responsiveness to anti-angiogenic therapy. Three of the SLC family members identified in NAXIVA responders were associated with favourable prognosis in TCGA data, where they were also predictive of immune microenvironment and drug response^32^. Considering other upregulated metabolic genes, *ALDOB* has also been reported to have prognostic significance in RCC^33^. These genes are expressed by normal renal tubules, so they may mark well-differentiated, less aggressive tumours.

A further question in the treatment of metastatic ccRCC is the potential synergistic effect of combining immunotherapy with VEGFR-directed TKIs, where the TKI is proposed to boost the effect of immunotherapy. Pre-clinical data indicates that VEGFR-TKIs enhance immunity by a variety of effects, including the reduction of immune suppressive myeloid cells in the tumour microenvironment (TME)^34–37^. In our data, we did not find any clear evidence of axitinib altering the immune profile and the overall immune phenotype remained stable on treatment. This is consistent with a study of neoadjuvant pazopanib in localised RCC, which did not find any change in immune signatures on treatment^20^. Axitinib has a narrow range of targets compared to other TKIs used in RCC^38^, so these observations do not rule out an immune modulatory effect of TKIs that target a wider range of receptors such as lenvatinib or cabozantanib.

Induction of PlGF at week 3 was a key marker of good outcome in NAXIVA. Biological heterogeneity, both between patients and within tumours, is a challenge for the development of baseline predictive biomarkers, whose limited performance could be surpassed by dynamic measurement of blood markers such as PlGF. Early blood biomarker changes may have significant clinical utility as they are readily assayed in the clinic. In the scenario of VTT management, it may give confidence in continuing with neoadjuvant therapy against proceeding directly to surgery. Our data do not provide a mechanism for the PlGF induction; however, we hypothesise that in responders, the effective blockade of the VEGFR axis induces a hypoxic response with increased production of VEGF-A and PlGF as a compensatory mechanism. It is not clear whether the responders and non-responders are biologically distinct in this respect, or whether there is a spectrum of effects depending on the degree of VEGFR inhibition achieved. PlGF is reported to potentiate the effectiveness of VEGF signalling, and so it may be a mechanism to overcome blockade^39^, particularly important in pathological angiogenesis compared to physiological angiogenesis^40^. PlGF is an attractive marker for further exploration as there is an existing clinical assay used for pre-eclampsia, which might be re-purposed.

A challenge in the analysis of small clinical trial datasets with extensive translational analysis is the large number of parameters assessed for predictive value in comparison to the number of patients enrolled in the study. ML approaches may enhance the analysis of similar datasets. The ML model based on baseline features achieved good performance for response prediction on an internal validation set. Performance is enhanced by data from week 3, again demonstrating the potential value of dynamic marker assessment. The models are limited by a small dataset and no current external validation data, but do provide an indication of putative response markers. The models selected a small number of factors based on plasma and tissue measurements which could be readily translated into the clinic. The result for CCL17 illustrates the utility of the ML approach, as this cytokine was assigned high priority by the ML models despite not being seen in our initial single parameter screens of the data. CCL17 is an important regulator of T cell immunity, acting on CCR4, and has been shown to be negatively prognostic in RCC^41^. The ML approach provided some insight into the interaction between the different features, particularly with responders being low in both IL-12p70 and CCL17. Changes in sTie-2 were also important in the dynamic model, which could be an alternative pathway for angiogenesis^42^; however, we interpret this finding with caution: although the fold change was consistent, the absolute changes in sTie-2 concentration in each patient were small. RNA-seq data was not included in our ML model due to the potential for noise amplification of adding several thousand differentially expressed genes to the other parameters. We observed differences in RNA and protein results; for instance, CCL17 was undetected in our RNA-seq and present at low levels in published single-cell datasets. This may suggest low transcript expression in tumours, making detection challenging, or indicate the importance of a non-tumour source, such as primary or secondary lymphoid tissue.

The study is limited by the small size of the trial, with only 20 participants, and by the lack of an external validation set for the key parameters identified, due to the unique nature of the study. We are restricted in both respects by the lower prevalence of VTT relative to all RCC cases; specific VTT management has been the subject of phase II trials to date, but a dedicated phase III VTT trial is likely unfeasible. Thus, we are limited to suggesting these markers as priorities for further work. Axitinib has been superseded by more active treatment combinations of TKIs and immunotherapy in the metastatic setting^43–45^, which is now being explored in pre-operative trials^13^; nonetheless, the TKI monotherapy in NAXIVA provides a useful comparator to any translational data arising from these IO-TKI studies.

Beyond phase II trials of current treatments, newer agents such as more potent TKIs or bispecific immunotherapies may have application in improving oncologic outcomes for VTT patients. This must be balanced against risks of toxicity. Our investigations of the microenvironment and blood features have identified predictive biomarkers that might be validated in these studies, either alone or as a combined assay. It will be interesting to see whether the key features identified by our study, which mainly divide the patients into ‘immune’ and ‘angiogenic’, are still valuable when combination treatment is used, or whether others emerge. It is critical that a range of translational analysis approaches, including tumour, blood, RNA and protein-based approaches linked to advanced cancer imaging, are built into future study designs to gain a full understanding of the mechanisms of tumour response and resistance.

## MATERIALS & METHODS

### Participants

NAXIVA was a single arm, single agent, phase II, open label, multicentre UK based study (NCT03494816, UK ethical approval REC reference: 17/EE/0240). Full study details including the trial protocol have been previously published^7^. Key inclusion criteria included: age >18, T3a, T3b or T3c, N0/N1, M0/1, biopsy proven clear cell RCC, suitable for immediate surgery. The baseline characteristics of the patients are summarised in the clinical publication^7^. Patients were treated with axitinib at a starting dose of 5mg BD, escalated to 7mg BD and then 10mg BD every 2 weeks. Drug was stopped a minimum of 36 hours and maximum of 7 days before surgery. The 20 evaluable patients in the intention to treat population in the main trial are included in the current study. Additional samples from ten untreated RCC patients with VTT were obtained from the ARTIST study (NCT04060537, UK ethical approval REC reference: 20/EE/0200). All patients were consented following GCP principles and the nature and possible consequences of the studies were explained. The studies were performed in accordance with the Declaration of Helsinki.

### Response evaluation

The technique for measuring VTT length by MRI is detailed in the original clinical report^7^; summarised as follows: Calculate the sum of (i) length of RV thrombus; (ii) the length of IVC tumour thrombus above the renal vein (measured from midpoint of the ostium of RV + IVC to tip of tumour thrombus); (iii) the length of IVC tumour thrombus below the renal vein (measured from midpoint of the ostium of RV + IVC to the tip of tumour thrombus). The percentage change in length at each timepoint (LT) compared to the length at baseline (LB) was calculated as (LT-LB)/LB*100.

### Histology & Image Analysis

Immunohistochemistry was performed on the Leica Bond III platform by standard automated procedure. The following antibodies were used: CD8 (4B11 Leica PA0183), CD31 (JC70A Leica PA0414), Ki67 (MIB-1Dako M7240). For immunofluorescence, 3-micron formalin-fixed paraffin-embedded (FFPE) sections were dewaxed in xylene and rehydrated in graded alcohols. Heat Induced Epitope Retrieval was performed in in Tris-EDTA pH9. After blocking, slides were incubated with primary antibodies at 4 °C overnight. Antibodies used were as follows: CD31 (JC/70A, Abcam), CD34 (AF7227, R&D Systems), SMA (ab5694, Abcam), CD68 (KP1, Invitrogen), Ki67 (EPR3610, Abcam), CD8 (SP16, Invitrogen), Granzyme B (NCL-L-GRAN-B, Leica), PD-1 (AF1086, R&D Systems), CD4 (EPR6855, Abcam), FOXP3 (236A/E7, Abcam), CA9 (AF2188, R&D Systems), CD3 (D7A6E, Cell Signalling Technology). Samples were washed and incubated in fluorescently conjugated secondary antibodies. Nuclei were counterstained with DAPI. Whole slides were scanned at ×40 magnification on the Zeiss Axio Scan Z1 system.

Image analysis was performed using HALO Software (Indica Labs). Tumour area was outlined manually for all slides. Slides with inadequate tissue quality for quantification were excluded from the analysis. Pre-defined analysis settings were applied to all slides for objective quantification. Analysis algorithms as follows: HighPlex FL v3.1.0, Object Colocalization FL v1.0, Area Quantification FL v2.1.5, Area Quantification v2.4.3, Multiplex IHC v3.1.4.

### Flow cytometr

PBMC samples collected during the trial were thawed and re-suspended in X-VIVO complete media (Lonza). Fc receptor block was used (Miltenyi). Cells were stained using standardised antibody panels (Supplementary Information). Viability was assessed by Zombie Aqua viability dye (Biolegend). Samples were run on a BD Symphony instrument. Appropriate single stain compensation bead controls were used. Data was analysed using FlowJo software.

### Cytokine arrays

Cytokine arrays were run by the Core Biochemical Assay Laboratory at the Cambridge Biomedical Research Centre, according to manufacturer’s instructions. The following kits were used from MesoScale Discovery: Human 10-plex Cytokine Panel 1 K15050D, Human 10-plex ProInflammatory Cytokine K15049D-2, Human 10-plex Chemokine Panel 1 K15047D, V-PLEX Angiogenesis Panel 1 Human Kit, K15050D. Plates were analysed on an MSD s600 instrument and results calculated by MSD Workbench software.

### Statistical analysis

Statistical analysis was conducted using R’s ggpubr package. For two-way comparisons, unpaired two-tailed student t-test was used with Bonferroni multiplicity correction where appropriate. For multiple comparisons, one-way ANOVA was used with Tukey’s post-hoc test. Correlation was by simple linear regression. All boxplots: centre line, median; box limits, upper and lower quartiles; whiskers, largest / smallest value or 1.5X interquartile range.

### Data availability

De-identified RNA-seq data will be made deposited in an appropriate repository on acceptance of the manuscript. De-identified imputed, normalised data set which was input to the machine learning models is available on request. Clinical data sharing will be shared where appropriate on correspondence with the author.

### Code availability

Machine learning code will be made available on Github on acceptance of the manuscript.

### Single cell analysis

To assess the expression of key genes in single cells derived from patients with RCC, we downloaded data from https://www.cell.com/cancer-cell/fulltext/S1535-6108(22)00548-7 via Mendeley Data: https://data.mendeley.com/datasets/g67bkbnhhg/1. To convert from.h5ad object to Seurat object we used sceasy (https://github.com/cellgeni/sceasy), prior to normalisation by mitochondrial content with SCTransform (from the R package Seurat_4.3.0). The average expression, and the percentage of cells that expressed the genes of interest was plotted. For clarity, we restricted cell types to endothelial, fibroblasts, and RCC cells as other cell types did not express the genes of interest (data not shown).

To plot the relative prevalence of cell types within different tissue compartments, we used code developed in https://www.cell.com/cancer-cell/fulltext/S1535-6108(22)00548-7 and documented in https://github.com/ruoyan-li/RCC-spatial-mapping. Briefly, we calculated the observed and expected number of cells of all cell types/subtype across different tissues. Adrenal metastasis and tumour thrombus were excluded from this analysis as they were only sampled in single patients. We also excluded blood cells as no endothelial, fibroblasts, and RCC cells were expected in this compartment.

### RNA-seq

#### RNA extraction & sequencing

RNA was extracted from formalin-fixed paraffin-embedded (FFPE) tissue using the ReliaPrep FFPE Total RNA kit, according to manufacturer’s instructions, and assessed by Qubit and Agilent RNA ScreenTape System and for quantity and quality. RNA library preparation was done using the Watchmaker Genomics RNA Library Prep Kit with Polaris Depletion, according to manufacturer’s instructions and running 18X PCR cycles for each sample. Indexing was done using the xGen™ Stubby Adaptor and UDI primers from Integrated DNA Technologies™, and sequencing via Illumina sequencing. The samples were run on an S4 flow cell on NovaSeq6000 with a read length of PE50. Manufacturer’s instructions followed for the run, including spike-in of 1% PhiX.

#### RNA-seq processing and analysis

Reads were mapped using Salmon (v1.10.0) with GRCh38.p44 from Gencode. Samples were only included in analysis if the sequencing duplication rate was < 65%. Genes were included if the maximum count per biopsy was > 10 and if more than 50% of the biopsies had counts > 0. Counts were normalised by variance-stabilised transformation and the top 500 genes generated principal components for the PCA plot. DESeq2 was used to identify differential expression between responders and non-responders, and between post-and pre-treatment samples. Genes satisfying P_adj_ < 0.05 and absolute Log_2_ Fold Change > 2 were used in pathway analysis via GO in the Cluster Profiler R package (v4.12.0) and plotted using EnrichPlot (v1.24.0). The same genes, and the genes for the features identified in the ML models, were highlighted on the published RNA-seq differential expression analysis data from IMmotion151^16^ to generate Figs. 6E and 8G, respectively.

#### RNA signature scores

The genes used for the Javelin Renal 101 Angio signature^15^: *NRARP, RAMP2, ARHGEF15, VIP, NRXN3, KDR, SMAD6, KCNAB1, CALCRL, NOTCH4, AQP1, RAMP3, TEK, FLT1, GATA2, CACNB2, ECSCR, GJA5, ENPP2, CASQ2, PTPRB, TBX2, ATP1A2, CD34, HEY2, EDNRB*. The genes used for the Javelin Renal 101 Immuno signature: *CD3G, CD3E, CD8B, THEMIS, TRAT1, GRAP2, CD247, CD2, CD96, PRF1, CD6, IL7R, ITK, GPR18, EOMES, SIT1, NLRC3, CD244, KLRD1, SH2D1A, CCL5, XCL2, CST7, GFI1, KCNA3, PSTPIP1*. The genes used for IMmotion151 Angio (C1/C2) signature^16^: *VEGFA, KDR, ESM1, PECAM1, ANGPTL4, CD34, FAP, FN1, COL5A1, COL5A2, POSTN, COL1A1, COL1A2, MMP2*. The genes used for IMmotion151 Immuno (C4/C5) signature: *CD8A, EOMES, PRF1, IFNG, CD274, CDK2, CDK4, CDK6, BUB1B, CCNE1, POLQ, AURKA, MKI67, CCNB2*. The genes used for the NAXIVA Angio signature: *PGF*, *TEK*, *PECAM1, CD34, VEGFA*. The genes used for the NAXIVA Immuno signature: *CCL17, IL12A, IL12B, IL7*. The counts were normalised by variance stabilised transformation, and the mean and standard deviation were calculated for each gene. For each patient, the score per gene is (expression – mean expression) / standard deviation across all patients. The total signature score per patient is the mean of the scores for each gene.

### Machine learning models

#### Training

We created a machine learning framework to predict response to axitinib. We used leave-one-out cross validation (LOOCV) on NAXIVA’s 20 patients to train and optimise the models. We used an increasing number of features, starting with baseline biopsy and blood features (62 features, Fig. 4a); then adding growth factor fold-change features at Week 3 of axitinib treatment (69 features, Fig. S10). For each combination we retrained the framework and derived a new model. The full list of features can be found in Supplementary Information. The models included recursive feature elimination using a logistic regression estimator, and the predictions were done by logistic regression with stochastic gradient descent, all coded in Python using scikit-learn version 1.4. Before entering the machine learning algorithm, all data underwent three pre-processing steps: iterative imputation, min-max standardisation and collinearity reduction. The estimator used for the iterative imputation was Bayesian Ridge. Collinearity reduction removed all features with a Spearman’s Rank correlation above 0.75, retaining one feature at random from the collinear group. The Spearman’s correlation between all features is visualised in Fig. S10. We used a five-fold cross validation setup to optimise model hyperparameters in the LOOCV training set, covering the hyperparameter ranges shown in Supplementary Information. The optimisation was based on a grid search in the hyperparameter space to optimise the area under the receiver operating characteristic curve. Once the optimal hyperparameters were found (Supplementary Information), we determined model parameters by re-fitting the model to the training set. To increase the robustness of the model with this dataset, LOOCV was done in 20 iterations, leaving one of the NAXIVA patients out at a time. The prediction for each left-out patient was done according to the model trained on the remaining 19 patients.

#### Feature importance

We evaluated feature importance in two different steps. First, we computed the frequency with which features were selected after the recursive feature elimination. We repeated the process for each of the LOOCV iterations, which means that features could be selected between 0 and a maximum of 20 times. Fig. 4e displays only features that were chosen at least eight out of twenty times in each cross-validation loop. Second, we computed the importance (i.e. weight) of each individual feature within the logistic regression algorithm. The weights were averaged across the iterations in which they were picked.

## Acknowledgments

The authors would like to thank the staff at the core facilities that were key to the success of this work: The Histopathology Core, Genomics Core, and Compliance & Biobanking Team at the CRUK Cambridge Institute, the Cancer & Molecular Diagnostics Laboratory at the Cambridge Biomedical Campus, The Human Research Tissue Bank and The Core Biochemical Assay Laboratory at Cambridge University Hospitals NHS Foundation Trust, and the NIHR Cambridge BRC Cell Phenotyping Hub. We would also like to thank the staff at the Scottish Clinical Trials Research Unit for their support in delivering the NAXIVA trial.

## Contributions

Conception and design: GDS, SJW, MCO, JOJ. Model development: IM, HP, RW. Acquisition of the data: JOJ, SJW, JB, FE, AE, FAG, TJM, LW, AYW, SU. Analysis and interpretation of the data: RW, HP, JOJ, JDS, MDR, SJW, MCO, GDS. Drafting of the manuscript: RW, HP, MCO, GDS, JOJ. Review and revision of the manuscript: RW, HP, IM, JB, FE, AE, FAG, TJM, MDR, JDS, SU, LW, AYW, SJW, MCO, GDS, JOJ. Statistical analysis: JOJ, RW, HP. Supervision: GDS, MCO, JJ.

