## Supplementary Information for "Angiogenic and Immune Predictors of Neoadjuvant Axitinib Response in Renal Cell Carcinoma with Venous Tumour Thrombus"

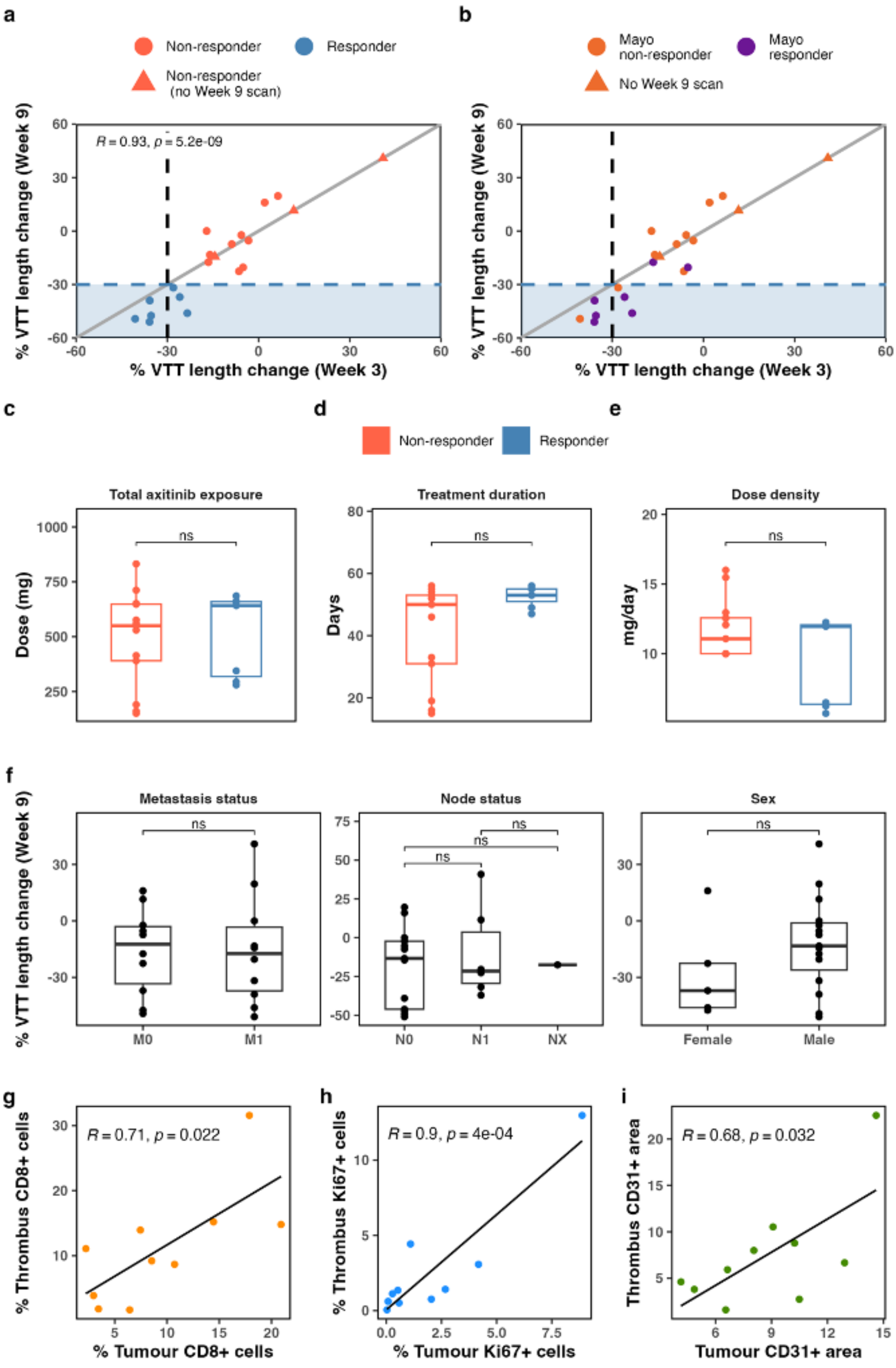


**Figure S1. Treatment response as assessed by VTT length on the NAXIVA Trial**
**a,** Association between MRI response at week 3 and week 9 (simple linear regression). Patients that stopped at week 3 are plotted along the line of week 3 = week 9 and were excluded from the

correlation analysis. **b,** Association between MRI response at week 3 and week 9, with response annotated by Mayo classification. Note that two patients with good length response would be classified as Mayo non-responders. **c,** Total axitinib received on trial. **d,** Duration of drug treatment on trial for each patient. **e,** Dose density (total mg / number of days treatment) for each patient **f,** VTT response by metastasis stage, lymph node stage and in male and female participants. **g-i,** Whole IHC slides were quantitated by HALO analysis for Ki67 **(g),** CD8 **(h)** and CD31 **(i)** in ten untreated paired primary tumours and VTTs.

(Pearson correlations between quantification of markers in VTT and the paired primary tumour are shown in **g-i**. For all other comparisons: unpaired Student's T-test with Bonferroni correction [ns: $p>0.05$, *: $p\leq0.05$, **: $p\leq0.01$, ***: $p\leq 0.001$])

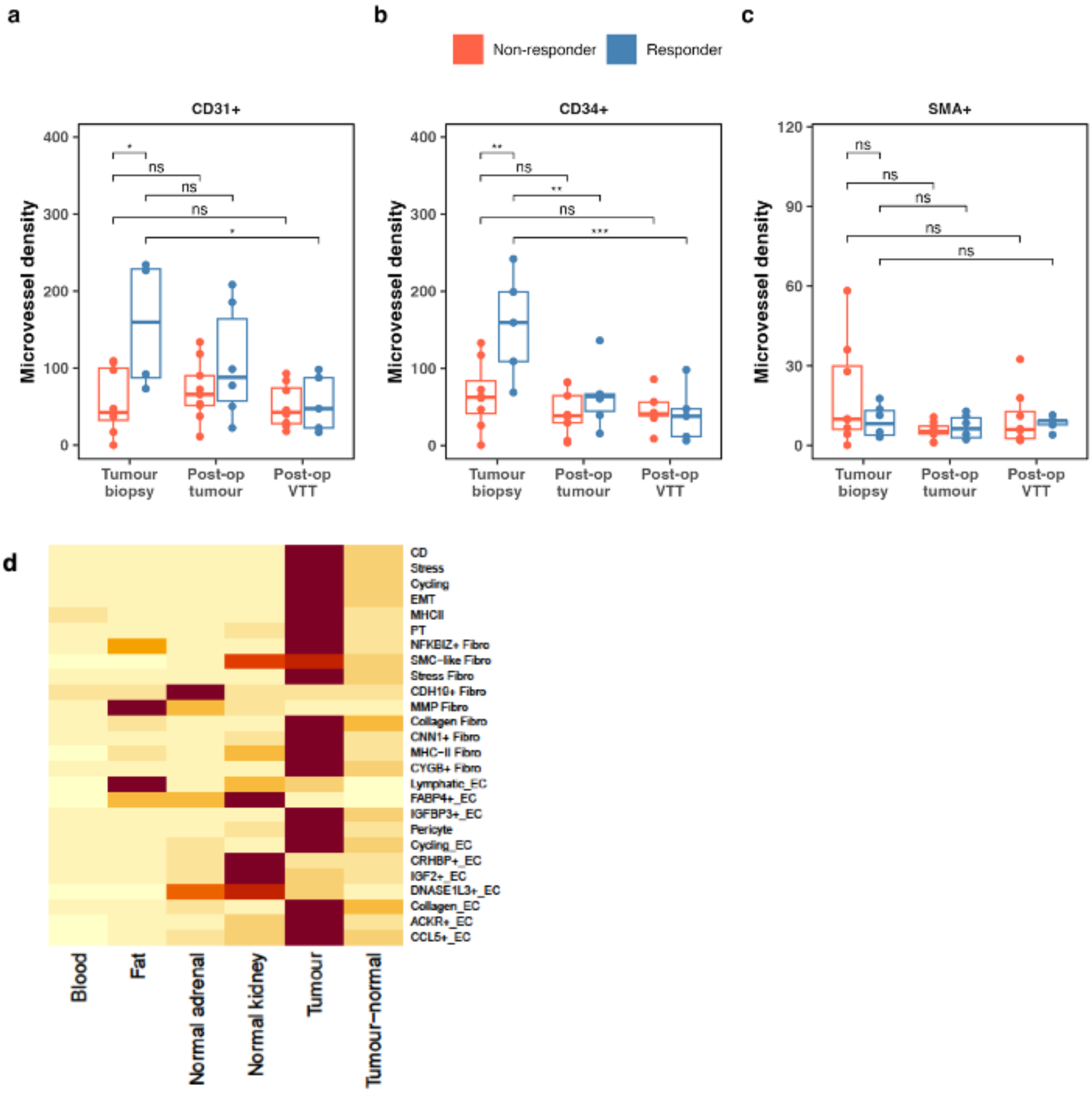


**Figure S2. Angiogenesis**

**a,** Change in CD31+ and **b,** CD34+ microvessel density and **c,** in SMA+ CAF area coverage (one-way ANOVA with Tukey's post-hoc test. [ns: p>0.05, *: p≤0.05, **: p≤0.01, ***: p≤ 0.001]). **d,** Cell prevalence by region in single cell dataset.

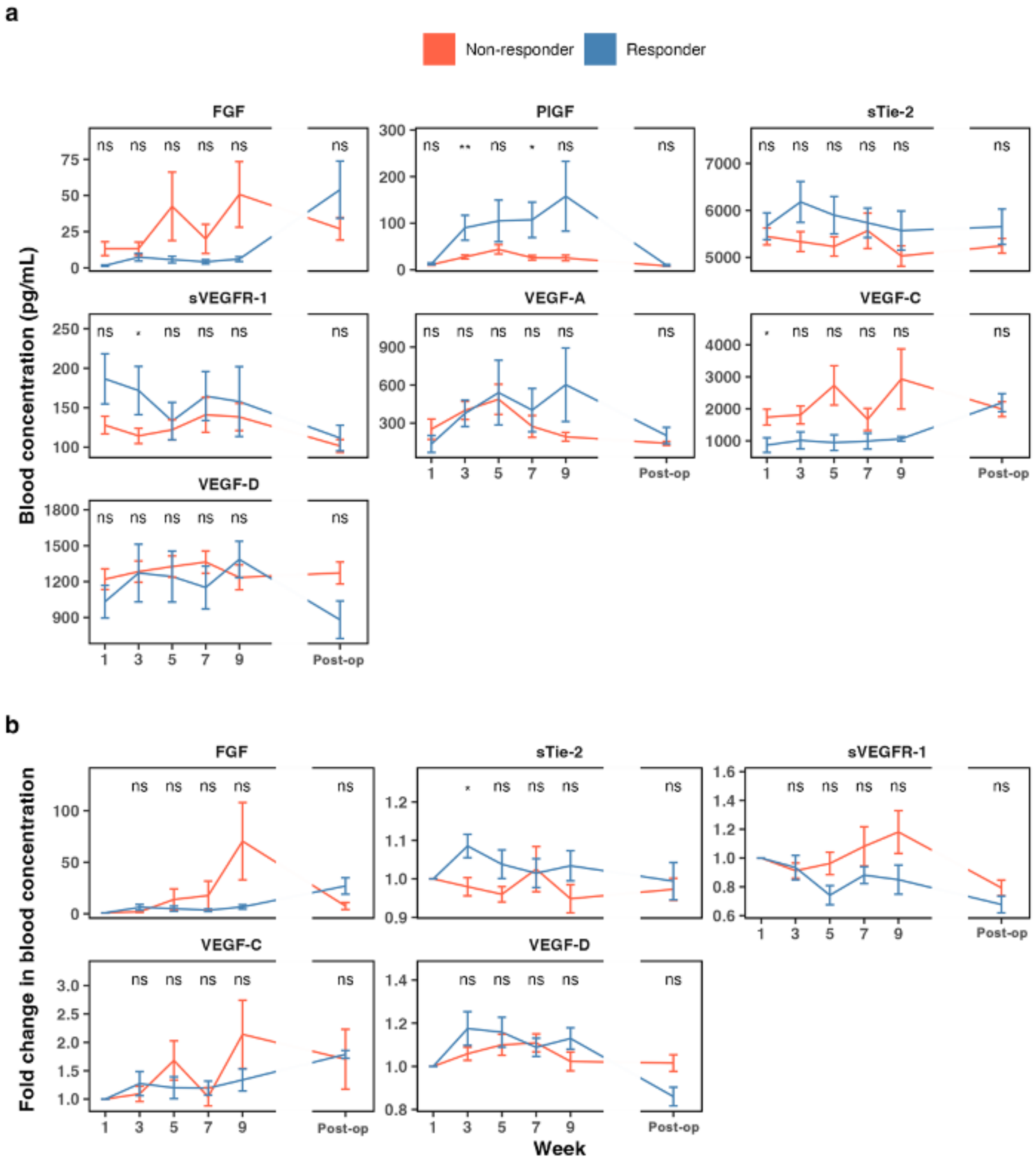


**Figure S3. Angiogenic factors**

**a,** Absolute and **b,** fold change measurements of angiogenic factors during the trial (unpaired Student's T-test [ns: p>0.05, *: p≤0.05, **: p≤0.01, ***: p≤ 0.001]).

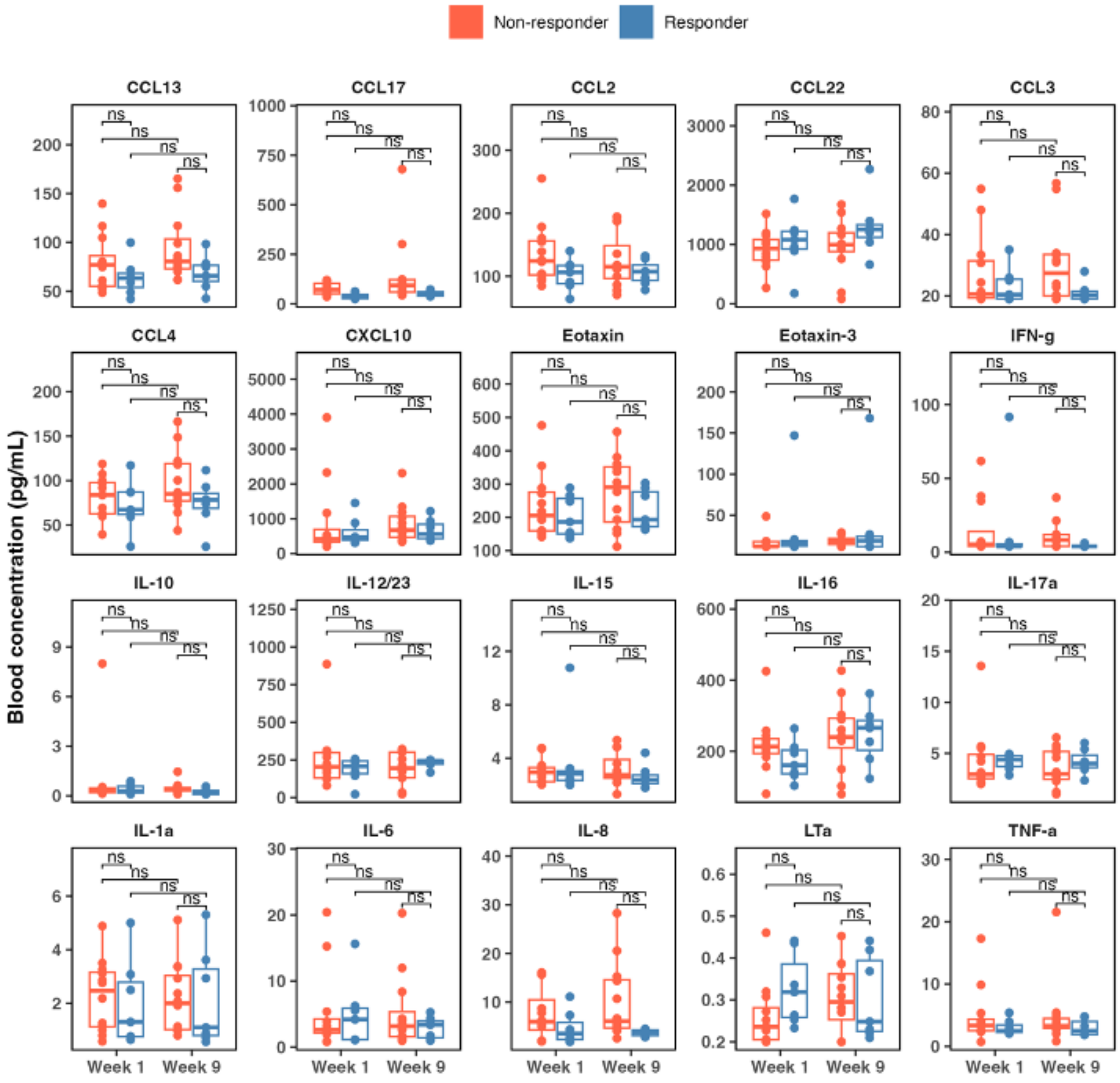


**Figure S4 Extended plasma cytokines**

Plasma cytokine levels were quantified at the beginning and end of axitinib treatment by cytokine array (one-way ANOVA with Tukey's post-hoc test [ns: $p>0.05$, *: $p\leq0.05$, **: $p\leq0.01$, ***: $p\leq 0.001$]).

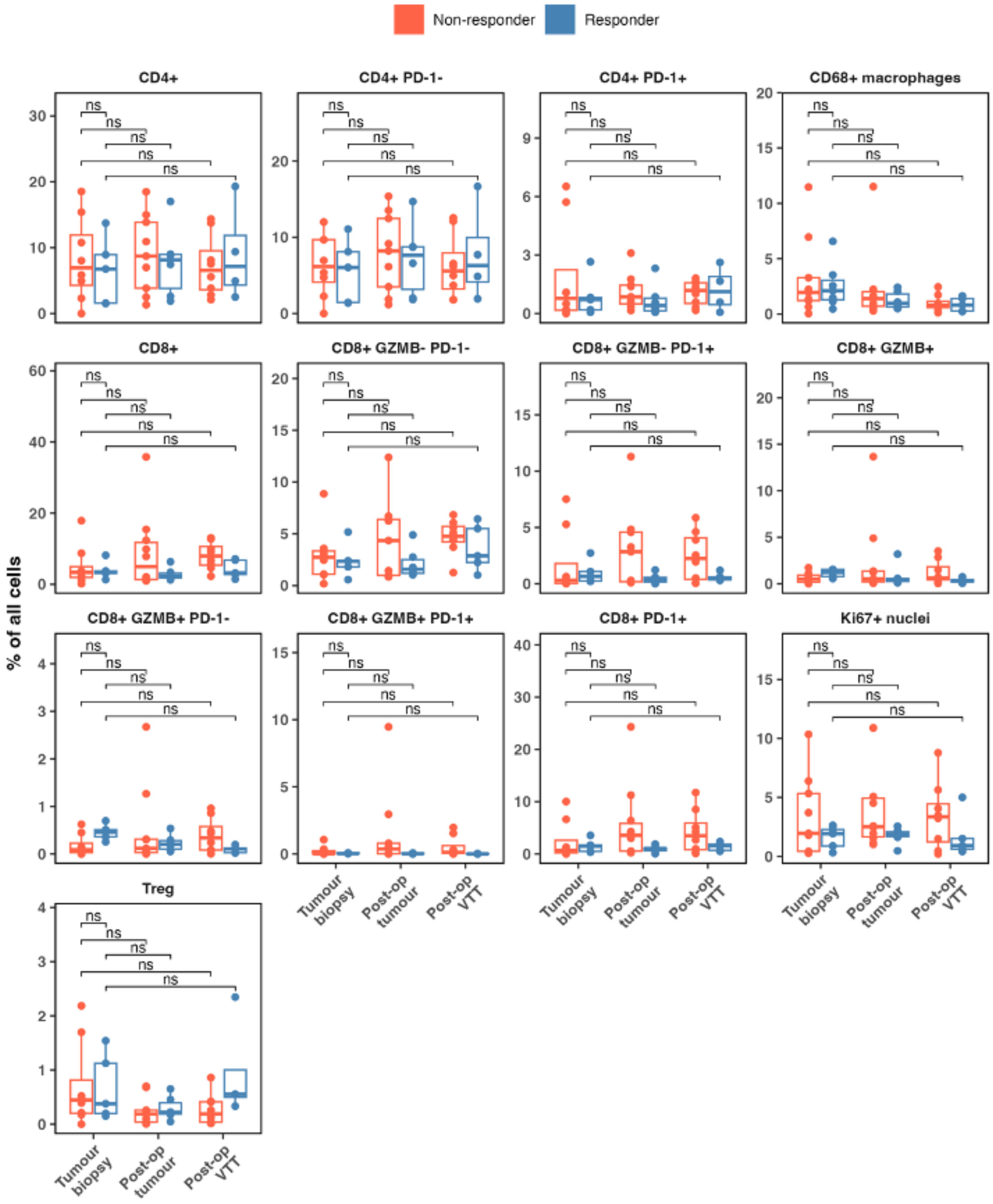


**Figure S5. Tissue immune subsets**
Multiplex immunofluorescence slides were quantified using HALO software.
(One-way ANOVA with Tukey's post-hoc test [ns: p>0.05, *: p≤0.05, **: p≤0.01, ***: p≤ 0.001])

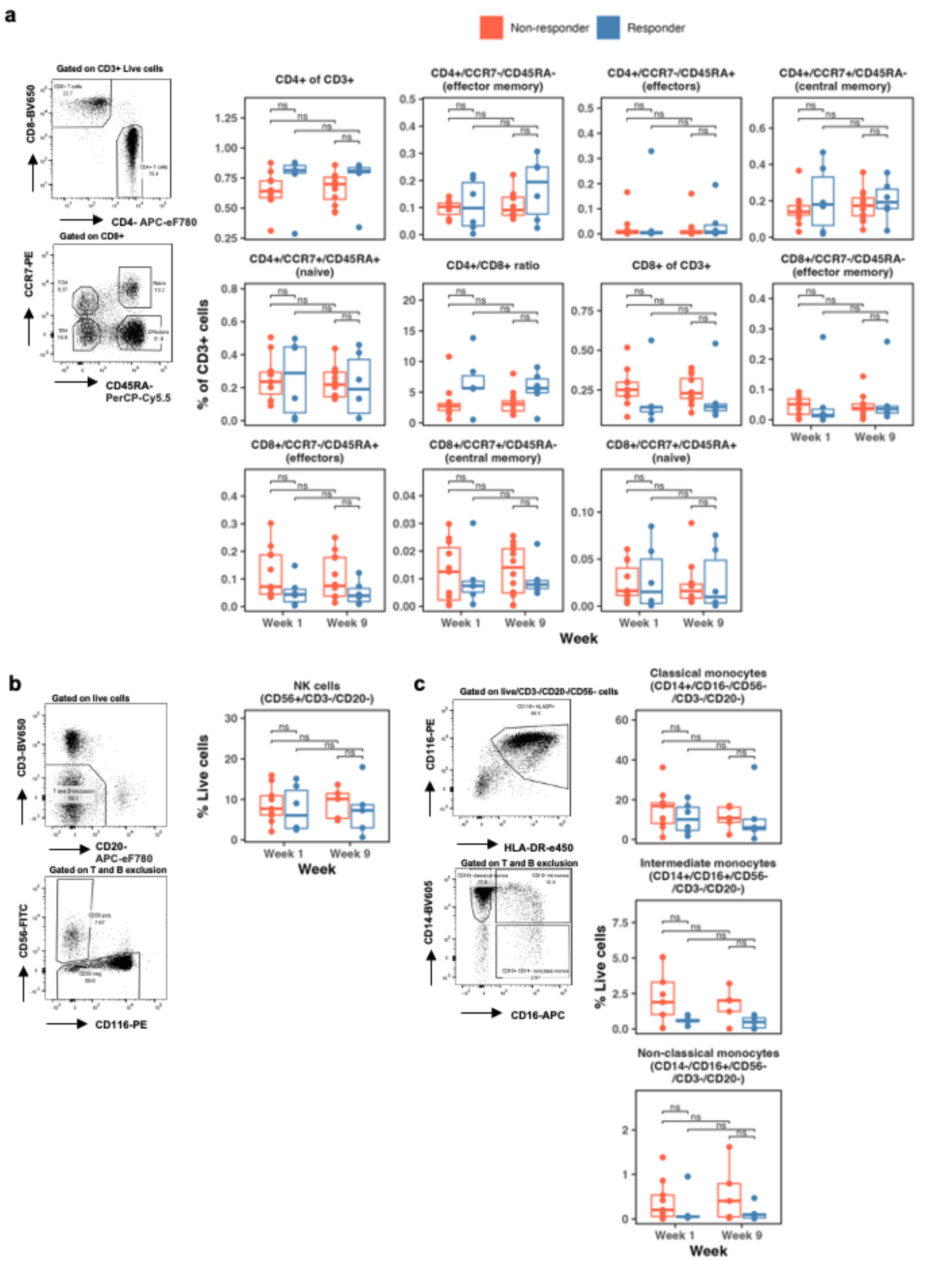


**Figure S6. Peripheral immune cell profile**

**a,** Representative image of gating strategy and CD4+ and CD8+ T cell subsets by CCR7 and CD45RA status, as percentage of CD3+/live cells (one-way ANOVA with Tukey's post-hoc test [ns: $p>0.05$, *: $p\leq0.05$, **: $p\leq0.01$, ***: $p\leq 0.001$]). **b-c,** Gating strategy & percentage of live cells for **b,** natural killer cells and **c,** monocyte subsets (unpaired Student's T-test with Bonferroni correction [ns: $p>0.05$, *: $p\leq0.05$, **: $p\leq0.01$, ***: $p\leq 0.001$]).

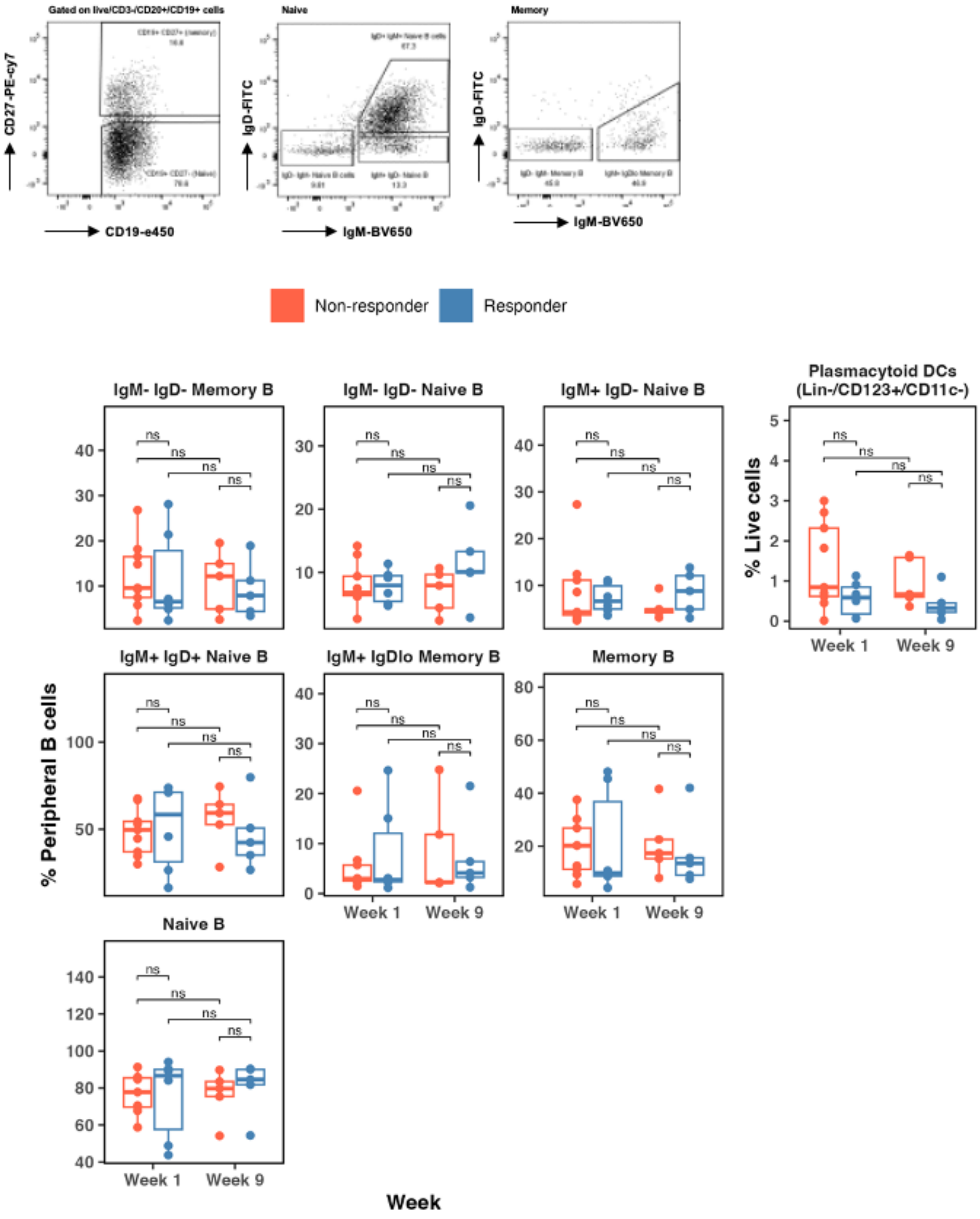


**Figure S7 Blood immune profile**
B cell subsets and plasmacytoid DCs from PBMCs taken at baseline and at the end of axitinib treatment were analysed by multicolour flow cytometry. B cell gating strategy is shown. (Unpaired Student's T-test with Bonferroni correction [ns: $p>0.05$, *: $p\leq0.05$, **: $p\leq0.01$, ***: $p\leq0.001$]).

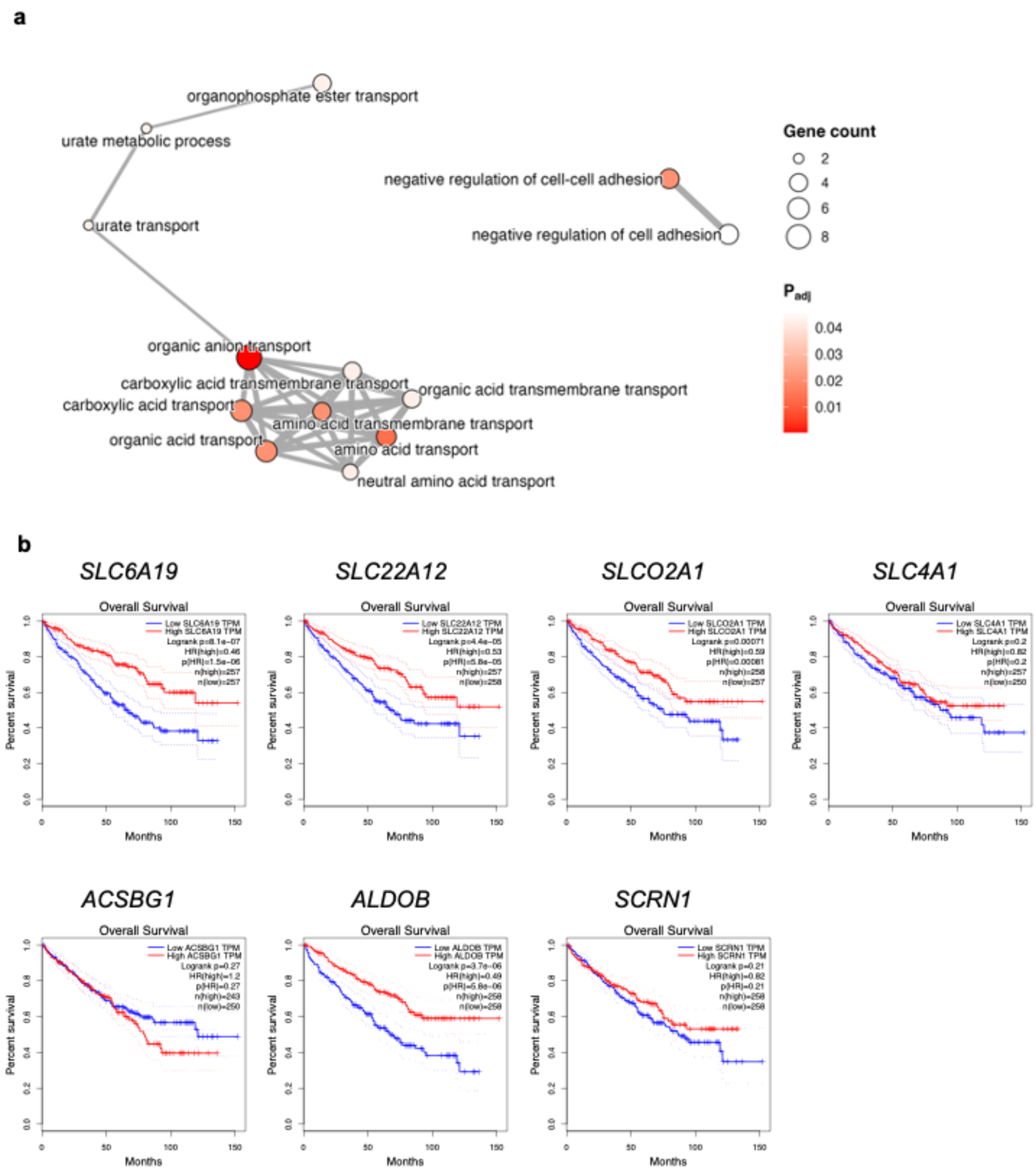


**Figure S8. RNA-seq GO analysis results**

**a,** GO analysis network for genes achieving P < 0.01 in the differential expression analysis. **b,** Survival plots of kidney cancer patients differentiated according to the expression of solute carrier genes highly expressed in responders to axitinib in NAXIVA.

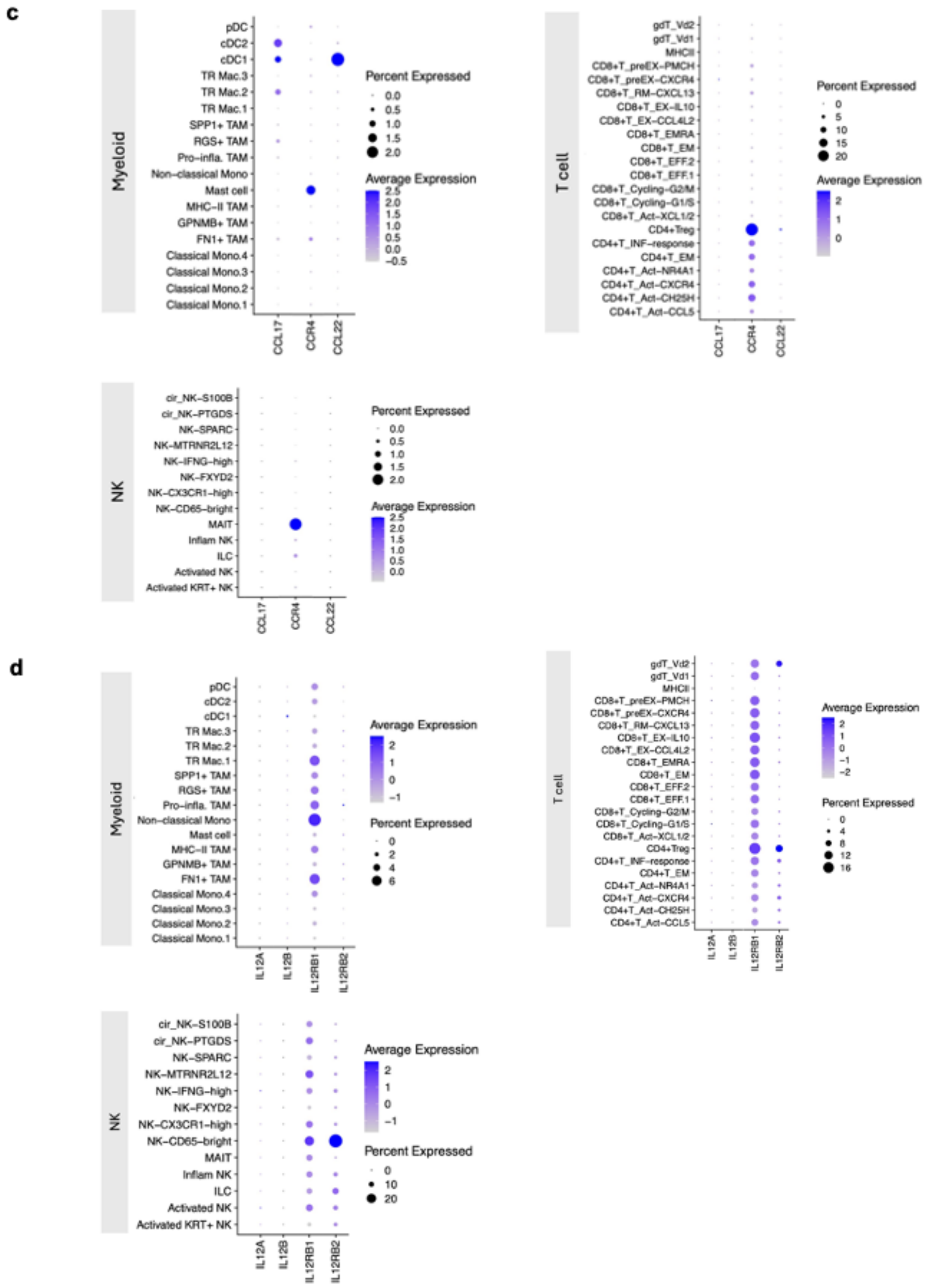


**Figure S9. scRNAseq compartments expressing key features**
**a,** Cells expressing CCL17, its receptor CCR4 and its antagonist CCL22 according to single cell RNA-seq data. **b,** Cells expressing IL12A, IL12B, and the receptors IL12R1 and IL12R2 according to single cell RNA-seq data.

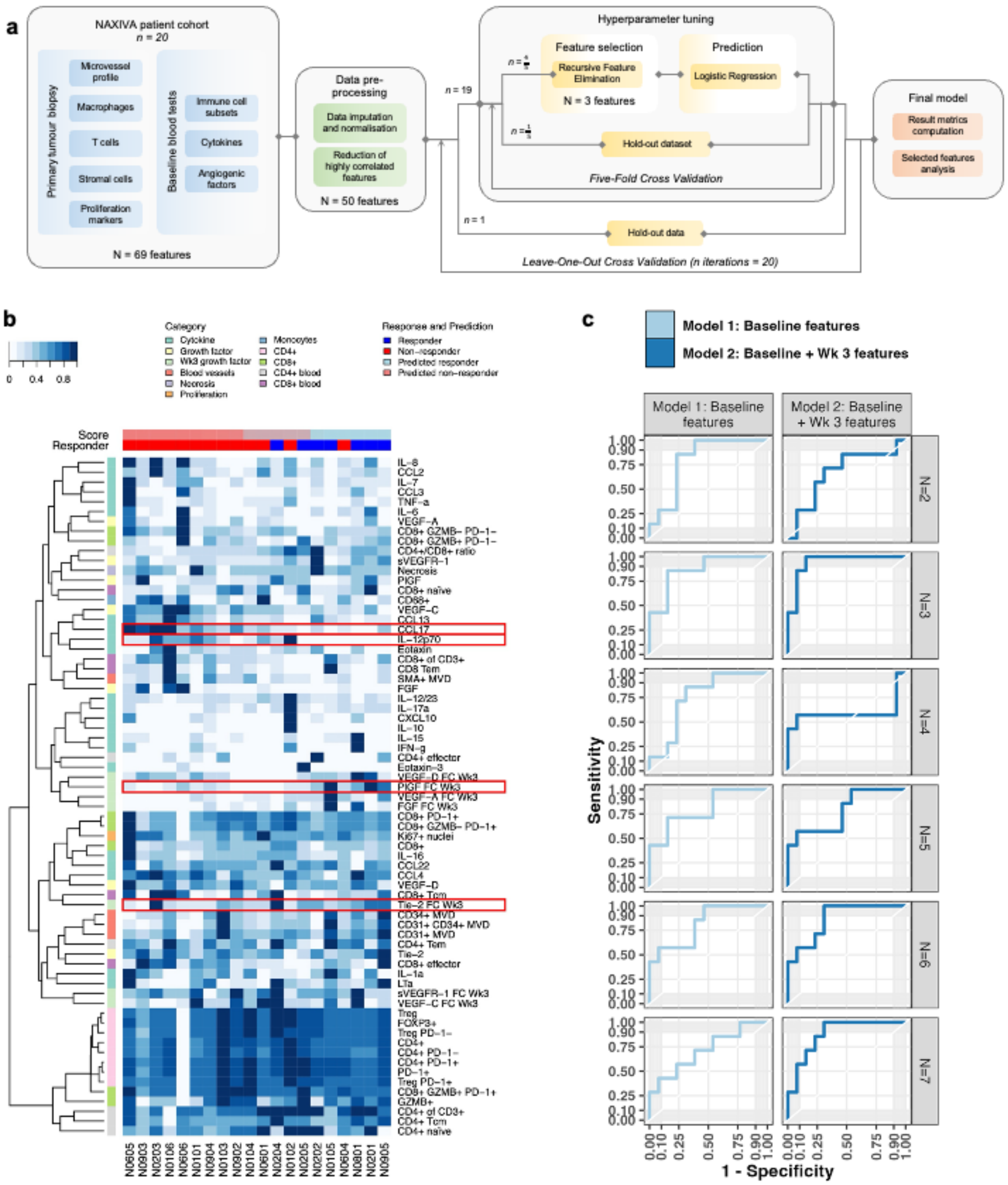


**Figure S10. Machine learning features and model design.**
**a,** Schematic of Model 2. **b,** Panel plot for Model 2 highlighting highest selected features. **c,** AUC plots for the optimisation of the number of features selected in the Logistic Regression RFE.

**Table S1. Flow cytometry panels**

| | FITC | PerCP-Cy5.5 | PE | PE-cy7 | APC | APC-eF780 | e450 | BV605 | BV650 | Zombie AQUA |
|---|---|---|---|---|---|---|---|---|---|---|
| T cell panel<br>*1 x 10^6 PBMC* | | CD45RA<br>eBioscience | CCR7<br>Biolegend | | | CD4<br>eBioscience | | CD3<br>Biolegend | CD8<br>Biolegend | LIVEDEAD<br>Biolegend |
| B cell panel<br>*1.5 x 10^6 PBMC* | IgD<br>BD | | | CD27<br>eBioscience | | CD20<br>eBioscience | CD19<br>BD | CD3<br>Biolegend | IgM<br>Biolegend | LIVEDEAD<br>Biolegend |
| Myeloid / NK panel<br>*1 x 10^6 PBMC* | CD56<br>eBioscience | CD123<br>eBioscience | CD116<br>eBioscience | CD11c<br>Miltenyi | CD16<br>eBioscience | CD20 & CD19<br>eBioscience | HLA-DR<br>eBioscience | CD14<br>Biolegend | CD3<br>Biolegend | LIVEDEAD<br>Biolegend |

**Table S2. Full list of features for machine learning models.**

| Measurement | Timepoint |
|---|---|
| VTT length | baseline |
| VTT length | week_3 |
| VTT length | week_9 |
| VTT fold reduction | week_3 |
| VTT fold reduction | week_9 |
| VTT absolute reduction | week_3 |
| VTT absolute reduction | week_9 |
| Mayo response | end |
| Sex | baseline |
| Node status | baseline |
| Metastasis status | baseline |
| Baseline grade | baseline |
| Surgery | surgery |
| Surgery grade | surgery |
| Necrosis | surgery |
| mg axitinib exposure | post_op |
| Days axitinib treatment | post_op |
| mg per day axitinib | post_op |
| VHL | baseline |
| BAP1 | baseline |
| PBRM1 | baseline |
| TERT | baseline |
| IFN-g | baseline |
| IL-10 | baseline |
| IL-12p70 | baseline |
| IL-6 | baseline |
| IL-8 | baseline |
| TNF-a | baseline |
| Eotaxin | baseline |
| Eotaxin-3 | baseline |
| CXCL10 | baseline |
| CCL2 | baseline |
| CCL13 | baseline |
| CCL22 | baseline |
| CCL3 | baseline |
| CCL4 | baseline |
| CCL17 | baseline |
| IL-1a | baseline |
| IL-12/23 | baseline |
| IL-15 | baseline |
| IL-16 | baseline |
| IL-17a | baseline |
| IL-7 | baseline |
| LTa | baseline |
| IFN-g | week_9 |
| IL-10 | week_9 |
| IL-12p70 | week_9 |
| IL-6 | week_9 |
| IL-8 | week_9 |
| TNF-a | week_9 |
| Eotaxin | week_9 |
| Eotaxin-3 | week_9 |
| CXCL10 | week_9 |
| CCL2 | week_9 |
| CCL13 | week_9 |
| CCL22 | week_9 |
| CCL3 | week_9 |
| CCL4 | week_9 |
| CCL17 | week_9 |
| IL-1a | week_9 |
| IL-12/23 | week_9 |
| IL-15 | week_9 |
| IL-16 | week_9 |
| IL-17a | week_9 |
| IL-7 | week_9 |
| LTa | week_9 |
| FGF | baseline |
| sVEGFR-1 | baseline |
| PlGF | baseline |
| Tie-2 | baseline |
| VEGF-A | baseline |
| VEGF-C | baseline |
| VEGF-D | baseline |
| FGF | week_3 |
| sVEGFR-1 | week_3 |
| PlGF | week_3 |
| Tie-2 | week_3 |
| VEGF-A | week_3 |
| VEGF-C | week_3 |
| VEGF-D | week_3 |
| FGF | week_5 |
| sVEGFR-1 | week_5 |
| PlGF | week_5 |
| Tie-2 | week_5 |
| VEGF-A | week_5 |
| VEGF-C | week_5 |
| VEGF-D | week_5 |
| FGF | week_7 |
| sVEGFR-1 | week_7 |

| | |
|---|---|
| PlGF | week_7 |
| Tie-2 | week_7 |
| VEGF-A | week_7 |
| VEGF-C | week_7 |
| VEGF-D | week_7 |
| FGF | week_9 |
| sVEGFR-1 | week_9 |
| PlGF | week_9 |
| Tie-2 | week_9 |
| VEGF-A | week_9 |
| VEGF-C | week_9 |
| VEGF-D | week_9 |
| FGF | post_op |
| sVEGFR-1 | post_op |
| PlGF | post_op |
| Tie-2 | post_op |
| VEGF-A | post_op |
| VEGF-C | post_op |
| VEGF-D | post_op |
| Necrosis | baseline |
| Necrosis | post_op_tumour |
| Necrosis | post_op_thrombus |
| CD31+ area | baseline |
| CD34+ area | baseline |
| CD31+ CD34+ area | baseline |
| CD31+ area | post_op_tumour |
| CD34+ area | post_op_tumour |
| CD31+ CD34+ area | post_op_tumour |
| CD31+ area | post_op_thrombus |
| CD34+ area | post_op_thrombus |
| CD31+ CD34+ area | post_op_thrombus |
| CD68+ | baseline |
| CD68+ | post_op_tumour |
| CD68+ | post_op_thrombus |
| SMA+ area | baseline |
| SMA+ area | post_op_tumour |
| SMA+ area | post_op_thrombus |
| Ki67+ nuclei | baseline |
| Ki67+ nuclei | post_op_tumour |
| Ki67+ nuclei | post_op_thrombus |
| CD8+ | baseline |
| GZMB+ | baseline |
| CD8+ PD-1+ | baseline |
| CD8+ GZMB+ PD-1- | baseline |
| CD8+ GZMB- PD-1+ | baseline |
| CD8+ GZMB+ PD-1+ | baseline |
| CD8+ GZMB- PD-1- | baseline |
| CD8+ | post_op_tumour |
| GZMB+ | post_op_tumour |
| CD8+ PD-1+ | post_op_tumour |
| CD8+ GZMB+ PD-1- | post_op_tumour |
| CD8+ GZMB- PD-1+ | post_op_tumour |
| CD8+ GZMB+ PD-1+ | post_op_tumour |
| CD8+ GZMB- PD-1- | post_op_tumour |
| CD8+ | post_op_thrombus |
| GZMB+ | post_op_thrombus |
| CD8+ PD-1+ | post_op_thrombus |
| CD8+ GZMB+ PD-1- | post_op_thrombus |
| CD8+ GZMB- PD-1+ | post_op_thrombus |
| CD8+ GZMB+ PD-1+ | post_op_thrombus |
| CD8+ GZMB- PD-1- | post_op_thrombus |
| CD4+ | baseline |
| FOXP3+ | baseline |
| PD-1+ | baseline |
| Treg | baseline |
| Treg PD-1+ | baseline |
| Treg PD-1- | baseline |
| CD4+ PD-1+ | baseline |
| CD4+ PD-1- | baseline |
| CD4+ | post_op_tumour |
| FOXP3+ | post_op_tumour |
| PD-1+ | post_op_tumour |
| Treg | post_op_tumour |
| Treg PD-1+ | post_op_tumour |
| Treg PD-1- | post_op_tumour |
| CD4+ PD1+ | post_op_tumour |
| CD4+ PD-1- | post_op_tumour |
| CD4+ | post_op_thrombus |
| FOXP3+ | post_op_thrombus |
| PD-1+ | post_op_thrombus |
| Treg | post_op_thrombus |
| Treg PD-1+ | post_op_thrombus |
| Treg PD-1- | post_op_thrombus |
| CD4+ PD1+ | post_op_thrombus |
| CD4+ PD-1- | post_op_thrombus |
| CD4+ of CD3+ | baseline |
| CD8+ of CD3+ | baseline |
| CD4+/CD8+ ratio | baseline |
| CD4+ effector | baseline |
| CD4+ naïve | baseline |
| CD4+ Tcm | baseline |
| CD4+ Tem | baseline |
| CD8+ effector | baseline |
| CD8+ naïve | baseline |

| | |
|---|---|
| CD8+ Tcm | baseline |
| CD8 Tem | baseline |
| CD4+ of CD3+ | post_op |
| CD8+ of CD3+ | post_op |
| CD4+/CD8+ ratio | post_op |
| CD4+ effector | post_op |
| CD4+ naïve | post_op |
| CD4+ Tcm | post_op |
| CD4+ Tem | post_op |
| CD8+ effector | post_op |
| CD8+ naïve | post_op |
| CD8+ Tcm | post_op |
| CD8 Tem | post_op |

**Table S3. Selected hyperparameters for each leave-one-out iteration of the model.**

| Model | Method | Algorithm | Iteration | hyperparams |
|---|---|---|---|---|
| Baseline | Logistic Regression RFE | Logistic SGD | 0 | [{'alpha': 0.0001, 'eta0': 0.01, 'learning_rate': 'optimal', 'penalty': 'l1'}] |
| Baseline | Logistic Regression RFE | Logistic SGD | 1 | [{'alpha': 0.0001, 'eta0': 0.01, 'learning_rate': 'optimal', 'penalty': 'l1'}] |
| Baseline | Logistic Regression RFE | Logistic SGD | 2 | [{'alpha': 0.0001, 'eta0': 0.01, 'learning_rate': 'optimal', 'penalty': 'l1'}] |
| Baseline | Logistic Regression RFE | Logistic SGD | 3 | [{'alpha': 0.0001, 'eta0': 0.01, 'learning_rate': 'optimal', 'penalty': 'l1'}] |
| Baseline | Logistic Regression RFE | Logistic SGD | 4 | [{'alpha': 0.0001, 'eta0': 0.01, 'learning_rate': 'optimal', 'penalty': 'l1'}] |
| Baseline | Logistic Regression RFE | Logistic SGD | 5 | [{'alpha': 0.0001, 'eta0': 0.01, 'learning_rate': 'optimal', 'penalty': 'l1'}] |
| Baseline | Logistic Regression RFE | Logistic SGD | 6 | [{'alpha': 0.0001, 'eta0': 0.01, 'learning_rate': 'optimal', 'penalty': 'l1'}] |
| Baseline | Logistic Regression RFE | Logistic SGD | 7 | [{'alpha': 0.0001, 'eta0': 0.01, 'learning_rate': 'optimal', 'penalty': 'l1'}] |
| Baseline | Logistic Regression RFE | Logistic SGD | 8 | [{'alpha': 0.0001, 'eta0': 0.01, 'learning_rate': 'optimal', 'penalty': 'l1'}] |
| Baseline | Logistic Regression RFE | Logistic SGD | 9 | [{'alpha': 0.0001, 'eta0': 0.01, 'learning_rate': 'optimal', 'penalty': 'l1'}] |
| Baseline | Logistic Regression RFE | Logistic SGD | 10 | [{'alpha': 0.0001, 'eta0': 0.01, 'learning_rate': 'optimal', 'penalty': 'l1'}] |
| Baseline | Logistic Regression RFE | Logistic SGD | 11 | [{'alpha': 0.0001, 'eta0': 0.01, 'learning_rate': 'optimal', 'penalty': 'l1'}] |
| Baseline | Logistic Regression RFE | Logistic SGD | 12 | [{'alpha': 0.0001, 'eta0': 0.01, 'learning_rate': 'optimal', 'penalty': 'l1'}] |
| Baseline | Logistic Regression RFE | Logistic SGD | 13 | [{'alpha': 0.0001, 'eta0': 0.01, 'learning_rate': 'optimal', 'penalty': 'l1'}] |
| Baseline | Logistic Regression RFE | Logistic SGD | 14 | [{'alpha': 100, 'eta0': 0.1, 'learning_rate': 'invscaling', 'penalty': 'l2'}] |
| Baseline | Logistic Regression RFE | Logistic SGD | 15 | [{'alpha': 0.0001, 'eta0': 0.01, 'learning_rate': 'optimal', 'penalty': 'l1'}] |
| Baseline | Logistic Regression RFE | Logistic SGD | 16 | [{'alpha': 100, 'eta0': 0.1, 'learning_rate': 'invscaling', 'penalty': 'l2'}] |
| Baseline | Logistic Regression RFE | Logistic SGD | 17 | [{'alpha': 0.0001, 'eta0': 0.01, 'learning_rate': 'optimal', 'penalty': 'elasticnet'}] |
| Baseline | Logistic Regression RFE | Logistic SGD | 18 | [{'alpha': 0.0001, 'eta0': 0.01, 'learning_rate': 'optimal', 'penalty': 'l1'}] |
| Baseline | Logistic Regression RFE | Logistic SGD | 19 | [{'alpha': 0.0001, 'eta0': 0.01, 'learning_rate': 'optimal', 'penalty': 'l1'}] |
| Baseline + Wk3 | Logistic Regression RFE | Logistic SGD | 0 | [{'alpha': 0.001, 'eta0': 0.01, 'learning_rate': 'optimal', 'penalty': 'elasticnet'}] |
| Baseline + Wk3 | Logistic Regression RFE | Logistic SGD | 1 | [{'alpha': 0.0001, 'eta0': 0.01, 'learning_rate': 'invscaling', 'penalty': 'l2'}] |
| Baseline + Wk3 | Logistic Regression RFE | Logistic SGD | 2 | [{'alpha': 0.0001, 'eta0': 0.1, 'learning_rate': 'optimal', 'penalty': 'l1'}] |
| Baseline + Wk3 | Logistic Regression RFE | Logistic SGD | 3 | [{'alpha': 100, 'eta0': 0.1, 'learning_rate': 'invscaling', 'penalty': 'l2'}] |

| | | | | |
|---|---|---|---|---|
| Baseline + Wk3 | Logistic Regression RFE | Logistic SGD | 4 | [{'alpha': 0.0001, 'eta0': 0.01, 'learning_rate': 'optimal', 'penalty': 'l1'}] |
| Baseline + Wk3 | Logistic Regression RFE | Logistic SGD | 5 | [{'alpha': 0.001, 'eta0': 0.01, 'learning_rate': 'optimal', 'penalty': 'l2'}] |
| Baseline + Wk3 | Logistic Regression RFE | Logistic SGD | 6 | [{'alpha': 0.0001, 'eta0': 0.01, 'learning_rate': 'optimal', 'penalty': 'l1'}] |
| Baseline + Wk3 | Logistic Regression RFE | Logistic SGD | 7 | [{'alpha': 0.0001, 'eta0': 0.01, 'learning_rate': 'optimal', 'penalty': 'l1'}] |
| Baseline + Wk3 | Logistic Regression RFE | Logistic SGD | 8 | [{'alpha': 0.0001, 'eta0': 0.01, 'learning_rate': 'optimal', 'penalty': 'elasticnet'}] |
| Baseline + Wk3 | Logistic Regression RFE | Logistic SGD | 9 | [{'alpha': 10, 'eta0': 1, 'learning_rate': 'invscaling', 'penalty': 'l2'}] |
| Baseline + Wk3 | Logistic Regression RFE | Logistic SGD | 10 | [{'alpha': 0.0001, 'eta0': 0.01, 'learning_rate': 'optimal', 'penalty': 'elasticnet'}] |
| Baseline + Wk3 | Logistic Regression RFE | Logistic SGD | 11 | [{'alpha': 0.0001, 'eta0': 0.01, 'learning_rate': 'invscaling', 'penalty': 'l2'}] |
| Baseline + Wk3 | Logistic Regression RFE | Logistic SGD | 12 | [{'alpha': 0.0001, 'eta0': 0.01, 'learning_rate': 'optimal', 'penalty': 'l1'}] |
| Baseline + Wk3 | Logistic Regression RFE | Logistic SGD | 13 | [{'alpha': 0.0001, 'eta0': 1, 'learning_rate': 'optimal', 'penalty': 'elasticnet'}] |
| Baseline + Wk3 | Logistic Regression RFE | Logistic SGD | 14 | [{'alpha': 0.0001, 'eta0': 0.01, 'learning_rate': 'optimal', 'penalty': 'l2'}] |
| Baseline + Wk3 | Logistic Regression RFE | Logistic SGD | 15 | [{'alpha': 0.0001, 'eta0': 0.01, 'learning_rate': 'optimal', 'penalty': 'l1'}] |
| Baseline + Wk3 | Logistic Regression RFE | Logistic SGD | 16 | [{'alpha': 0.0001, 'eta0': 0.01, 'learning_rate': 'optimal', 'penalty': 'l1'}] |
| Baseline + Wk3 | Logistic Regression RFE | Logistic SGD | 17 | [{'alpha': 0.0001, 'eta0': 0.01, 'learning_rate': 'optimal', 'penalty': 'l1'}] |
| Baseline + Wk3 | Logistic Regression RFE | Logistic SGD | 18 | [{'alpha': 0.001, 'eta0': 0.1, 'learning_rate': 'optimal', 'penalty': 'l2'}] |
| Baseline + Wk3 | Logistic Regression RFE | Logistic SGD | 19 | [{'alpha': 0.0001, 'eta0': 0.01, 'learning_rate': 'optimal', 'penalty': 'l1'}] |

**Table S4. Hyperparameter ranges**

| Hyperparameter | Range |
|---|---|
| alpha | [0.0001, 0.001, 0.01, 0.1, 1, 10, 100] |
| penalty | ['l1', 'l2', 'elasticnet'] |
| learning_rate | ['optimal', 'invscaling', 'adaptive'] |
| eta0 | [0.01, 0.1, 1] |
